# Initial ciliary assembly in *Chlamydomonas* requires Arp2/3 complex-dependent endocytosis

**DOI:** 10.1101/2020.11.24.396002

**Authors:** Brae M Bigge, Nicholas E Rosenthal, Prachee Avasthi

## Abstract

Ciliary assembly, trafficking, and regulation are dependent on microtubules, but the mechanisms of ciliary assembly also require the actin cytoskeleton. Here, we dissect subcellular roles of actin in ciliogenesis by focusing on actin networks nucleated by the Arp2/3 complex in the powerful ciliary model, *Chlamydomonas*. We find the Arp2/3 complex is required for the initial stages of ciliary assembly when protein and membrane are in high demand but cannot yet be supplied from the Golgi complex. We provide evidence for Arp2/3 complex-dependent endocytosis of ciliary proteins, an increase in endocytic activity upon induction of ciliary growth, and relocalization of plasma membrane proteins to newly formed cilia. Our data support a new model of ciliary protein and membrane trafficking during early ciliogenesis whereby proteins previously targeted to the plasma membrane are reclaimed by Arp2/3 complex-dependent endocytosis for initial ciliary assembly.

**SUMMARY:** Using the ciliary model system *Chlamydomonas*, we find Arp2/3 complex-mediated endocytosis is needed to reclaim cell body plasma membrane for early ciliary assembly.

## INTRODUCTION

The cilium of the unicellular, green alga *Chlamydomonas reinhardtii* has long been used as a model due to its structural and mechanistic conservation relative to mammalian cilia. Cilia consist of microtubules that extend from the cell surface and are ensheathed in plasma membrane. Their assembly relies on microtubule dynamics and trafficking of protein and membrane (Nachury et al., 2010), as well as intraflagellar transport (IFT), a motor-based transport system that moves tubulin and other cargo through the cilium (Pedersen and Rosenbaum, 2008).

Although cilia are composed of microtubules and depend on them for assembly, the mechanisms governing ciliary maintenance and assembly extend to other cytoskeletal proteins, like actin. The microtubule organizing center, the centrosome, from which cilia are nucleated functions as an actin organizer (Farina et al., 2016; Inoue et al., 2019). In mammalian cells, cortical actin disruption results in increased ciliary length and percentage of ciliated cells (Kim et al., 2010; Park et al., 2008), and when ciliogenesis is triggered by serum starvation, preciliary vesicles are trafficked to the centriole where they fuse to form a ciliary vesicle around the budding cilium. In intracellular ciliogenesis, when Arp2/3 complex-branched actin is lost, vesicle fusion defects lead to depletion of preciliary vesicles at the centriole, suggesting a role for branched actin in intracellular ciliogenesis (Wu et al., 2018). Further, actin itself has been found within cilia, suggesting that actin is a key protein in ciliary maintenance and assembly (Kiesel et al., 2020).

*Chlamydomonas* cells are ideal for tackling the question of actin-dependent ciliary trafficking due to their lack of a cortical actin network and their ability to undergo consistent and robust ciliogenesis without requiring serum starvation. In *Chlamydomonas*, disruption of actin networks with Cytochalasin D resulted in short cilia (Dentler and Adams, 1992) and disruption with Latrunculin B (LatB), which sequesters monomers leading to filament depolymerization, resulted in short cilia and impaired regeneration (Avasthi et al., 2014; Jack et al., 2019). *Chlamydomonas* actin networks are required for accumulation of IFT machinery at the base of cilia and for entry of IFT material into cilia (Avasthi et al., 2014), as well as for trafficking of post-Golgi vesicles to cilia, synthesis of ciliary proteins, and organization of the ciliary gating region (Jack et al., 2019). Many advances in our understanding of the relationship between cilia and actin were discovered using *Chlamydomonas*.

The actin cytoskeleton of *Chlamydomonas* contains two actin genes: *IDA5*, a conventional actin with 91% sequence identity to human β-actin; and *NAP1*, a divergent actin that shares only 63% of its sequence with human β-actin (Hirono et al., 2003; Kato-Minoura et al., 1998). We consider NAP1 to be an actin-like protein as opposed to an actin related protein (ARP) because it has a higher sequency identity to actin than to conventional ARPs and because it is able to functionally compensate for the conventional IDA5 (Jack et al., 2019; Onishi et al., 2019, 2018, 2016). Under normal, vegetative conditions, IDA5 is the primary actin expressed, but when cells are treated with LatB, the LatB-insensitive NAP1 is upregulated (Hirono et al., 2003; Onishi et al., 2018, 2016). This separability of the two actins led to the discovery that they can compensate for each other in ciliary maintenance and assembly (Jack et al., 2019). Studies of actin’s role in ciliary assembly used global disruption, knocking out either one of the filamentous actins or acutely knocking out both, yet actin networks have diverse compositions and topologies that lead to specific subfunctions within cells.

Actin networks rely on actin binding proteins that contribute to the formation, arrangement, and function of the networks. One such actin binding protein is the Arp2/3 complex, which nucleates branched or dendritic networks. These networks are often involved in membrane remodeling functions, like lamellipodia and endocytosis (Campellone and Welch, 2010). The Arp2/3 complex consists of 7 subunits: Arp2, Arp3, and ARPC1-5 (**Figure S1**). Each subunit plays a specific role of varying importance in the nucleation process. ARPC2 and ARPC4 form the complex core and the primary contacts with the mother filament, Arp2 and Arp3 serve as the first subunits of the daughter filament, and ARPC1 and ARPC3 play a role in nucleation but are not critical for branch formation (Gournier et al., 2001; Robinson et al., 2001). Each of these subunits is found in *Chlamydomonas*, but they have a range of sequence homologies compared to conventional Arp2/3 complexes (**Figure S1**). The ARPC5 subunit has yet to be found in *Chlamydomonas*. ARPC5 is thought to be important for the association of ARPC1 to the complex, but a mammalian complex lacking ARPC5 and ARPC1 maintains some nucleating and branching activity and is able to cross-link actin (Gournier et al., 2001).

Here, using the chemical inhibitor CK-666 to inhibit the nucleating function of the Arp2/3 complex (Hetrick et al., 2013) and a genetic mutant of a critical Arp2/3 complex member, ARPC4 (Cheng et al., 2017; Li et al., 2019), we take a more delicate approach to investigating the actin’s in ciliary assembly by separating different actin networks into their subfunctions based on topology. Specifically, we probe the involvement of actin networks nucleated by the Arp2/3 complex in ciliary maintenance and assembly. This approach in these cells has allowed us to propose a new model implicating a subset of filamentous actin in redistribution of membrane and proteins for the initial stages of ciliogenesis.

## RESULTS

### Loss of Arp2/3 complex function inhibits normal regeneration and maintenance of cilia

To investigate the role of Arp2/3 complex-mediated actin networks in ciliary assembly, we used two tools. First, we used the chemical inhibitor CK-666 which blocks the nucleating ability of the Arp2/3 complex by binding the interface between Arp2 and Arp3 and locking the complex in an inactive state (Hetrick et al., 2013). The Arp2 and Arp3 subunits of the *Chlamydomonas* Arp2/3 complex are 75.1% and 61.5% similar to mammalian Arp2/3 complex respectively (**Figure S1**). This, along with the ability of CK-666 to recapitulate the effects of a genetic mutant of the Arp2/3 complex, suggest that CK-666 can target *Chlamydomonas* Arp2/3 complex. Second, we obtained a loss of function mutant of the critical Arp2/3 complex member, ARPC4 from the *Chlamydomonas* Resource Center (Cheng et al., 2017; Li et al., 2019). The *arpc4* mutant was confirmed via PCR and further evaluated using a genetic rescue where a V5-tagged ARPC4 construct is expressed in *arpc4* mutant cells, *arpc4:*ARPC4-V5. This was confirmed via PCR, western blot, and immunofluorescence (**Figure S1C-E**). While we can confirm the presence of ARPC4-V5 with immunofluorescence, the actual localization of ARPC4-V5 is not discernable due to diffuse signal (**Figure S1E**). This could be because all ARPC4-V5 in the cell is not being incorporated into active Arp2/3 complexes.

We probed the requirement for the Arp2/3 complex in maintenance of cilia by treating cells with CK-666 or the inactive control CK-689 for 2 hours until cilia reached a new steady state length. Consistent with previous results (Avasthi et al., 2014), CK-666 decreased ciliary length (**Figure 1A**). We saw no changes with the inactive CK-689 (**Figure 1A**) or when *arpc4* mutant cells lacking a functional Arp2/3 complex were treated with CK-666 (**Figure 1A**). Untreated *arpc4* mutant cells recapitulate the CK-666 result, showing decreased ciliary length when compared to wild-type cells (**Figure 1B**). This defect in ciliary length was rescuable with ARPC4-V5 (**Figure 1B**). Overall, this suggests the Arp2/3 complex is required for normal ciliary maintenance.

**Figure 1.**
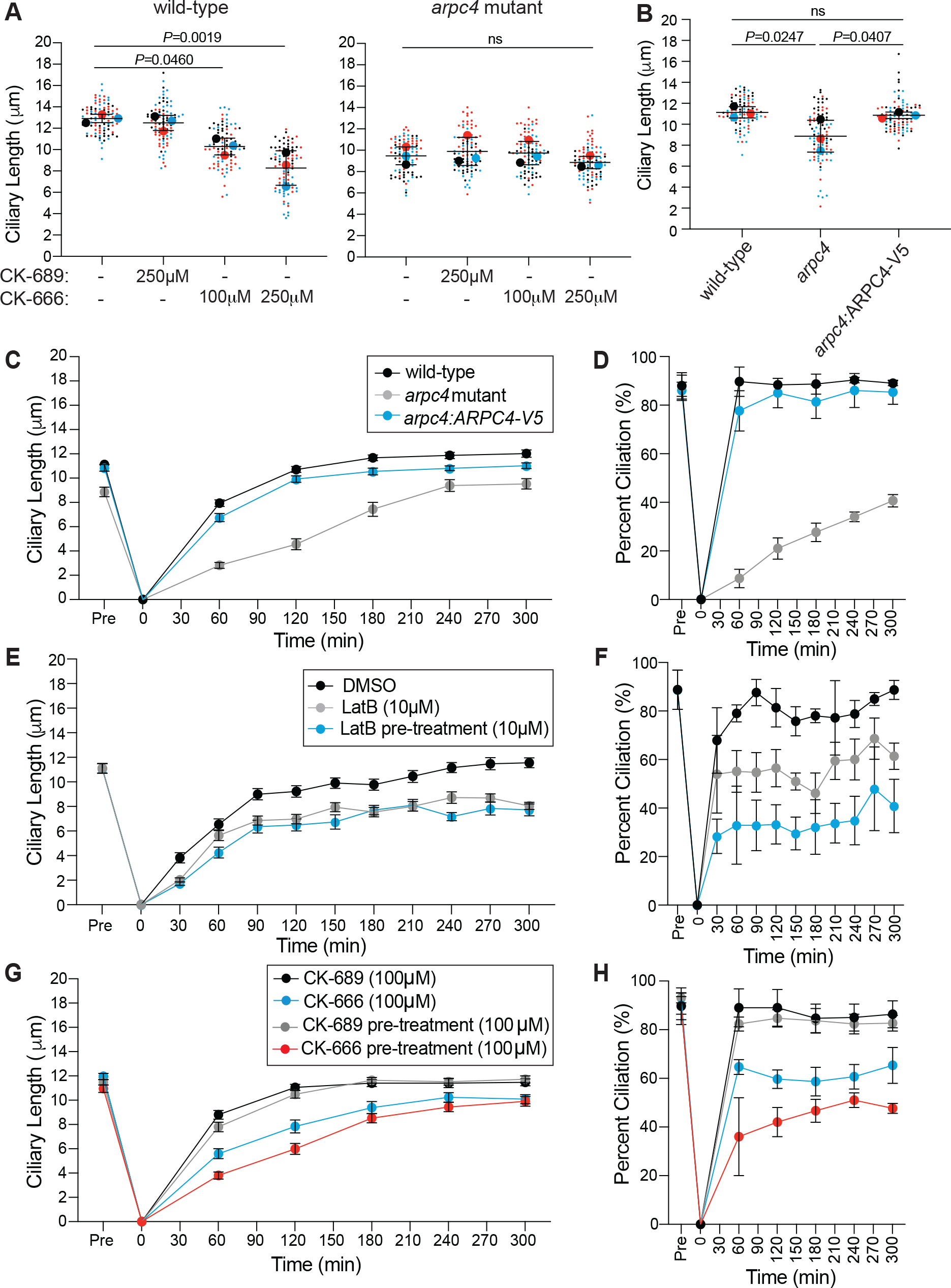
The Arp2/3 complex is required for normal ciliary maintenance and assembly. **A)** Wild-type and *arpc4* mutant cells were treated with 100μM or 250μM CK-666 or the inactive CK-689 for 2 hours. Superplots show the mean ciliary lengths from 3 separate biological replicates with error bars representing standard deviation. n=30 for each treatment in each biological replicate. **B)** Wild-type cells, *arpc4* mutant cells, and *arpc4:ARPC4-V5* cells steady state cilia were also measured with no treatment. Superplots show the mean of 3 biological replicates with error bars representing standard deviation. n=30 for each strain in each biological replicate. **C-D)** Wild-type cells and *arpc4* mutant cells were deciliated using a pH shock, cilia were allowed to regrow and ciliary length (C) and percent ciliation (D) were determined. Means are displayed with error bars representing 95% confidence interval (C) or standard deviation (D). n=30 (C) or n=100 (D) for each strain and each time-point in 3 separate biological replicates. For every time point except 0 min, P<0.0001 for both length and percent ciliation. **E-F)** *nap1* mutant cells were pre-treated with 10μM LatB for 30 minutes before deciliation or treated with LatB upon the return to neutral pH following deciliation. Ciliary length (E) and percent ciliation (F) were determined for each time point. Error bars represent 95% confidence interval (E) or standard deviation (F). n=30 (E) or n=100 (F) for 3 separate experiments. For every time point P>0.0001 between DMSO and treated samples, except 30min (10μM LatB) which is ns. **G-H)** Wild-type cells were pre-treated with CK-666 or the inactive CK-689 (100μM) for 1 hour before deciliation of treated with CK-666 or the inactive CK-689 (100μM) following deciliation. Ciliary length (G) and percent ciliation were measured (H). Error bars represent 95% confidence interval (G) or standard deviation (H). n=30 (G) or n=100 (H) for 3 separate experiments.

Next, we explored the involvement of the Arp2/3 complex in ciliary assembly where protein and membrane both from existing pools and from synthesis are in high demand (Diener et al., 2015; Jack et al., 2019; Nachury et al., 2010; Rohatgi and Snell, 2010; Wingfield et al., 2017). Cells were deciliated by low pH shock and allowed to synchronously regenerate cilia at normal pH (Lefebvre, 1995). *arpc4* mutant cells were slow to regenerate cilia, and roughly 60% of cells did not regrow cilia (**Figure 1C-D**). This phenotype was rescued by with ARPC4-V5 (**Figure 1C-D**). Importantly, the most severe defect in assembly is in initial steps when existing protein and membrane are being incorporated into cilia.

The striking decrease in ciliary assembly is puzzling because loss of the Arp2/3 complex, and therefore only a subset of actin filaments, results in a more dramatic phenotype than *nap1* mutants treated with LatB, which lack all filamentous actins (Jack et al., 2019). However, in *arpc4* mutant cells, a functional Arp2/3 complex never exists. In *nap1* mutant cells treated with LatB, treatment begins shortly after deciliation resulting in an acute perturbation. Further, LatB functions by sequestering actin monomers to promote filament disassembly, and the effects may not be immediate (Spector et al., 1989). Thus, there is likely a brief window where actin filaments can assert their initial role in ciliary regeneration before being depolymerized. To avoid this, we began LatB treatment in *nap1* mutants 30 minutes before deciliation, which allowed us to observe what happens when actin is not present immediately after deciliation. We see slightly decreased ciliary length consistent with the acute treatment but dramatically decreased percent ciliation, which is consistent with the *arpc4* mutant results (**Figure 1E-F**).

Treatment with CK-666, an Arp2/3 complex inhibitor, gives a similar result. In cells treated with CK-666 immediately following deciliation, the Arp2/3 complex may assert its role in assembly before being inhibited by CK-666. By pre-treating cells with CK-666 for 1 hour before deciliation, we can observe what happens when Arp2/3 complex function is lost immediately following deciliation. When we do so, we see a more dramatic defect in ciliary length and percent ciliation than we do with just acute CK-666 treatment (**Figure 1G-H**), suggesting the Arp2/3 complex is required for a very early initial step of assembly that occurs before we have a chance to treat the cells.

### The Arp2/3 complex is required for the incorporation of existing membrane and proteins for ciliary assembly

There are several actin-dependent steps of ciliary assembly, including incorporation of existing protein and membrane and synthesis and trafficking of new protein for cilia. Using a method to label nascent peptides, we found that loss of ARPC4 did not prevent upregulation of translation following deciliation (**Figure S3**). In this experiment, we halt translation and fluorescently label newly translated polypeptides. Wild-type and *arpc4* mutant cells were tested using this reaction either before deciliation, following deciliation and one hour of regrowth, or following deciliation and one hour of regrowth in cycloheximide (CHX), which inhibits protein synthesis by blocking the elongation step of protein translation. Wild-type and *arpc4* mutant cells displayed an increase in cell fluorescence following deciliation, especially around the nucleus, indicating an increase in protein synthesis following deciliation (**Figure S3**). Importantly, this increase in cell fluorescence was not significantly different between wild-type and *arpc4* mutant cells, suggesting loss of Arp2/3 complex function does not prevent upregulation of protein synthesis following deciliation. This also indicates *arpc4* mutant cells are aware their cilia were lost, as they respond with increased protein synthesis.

Given that *arpc4* mutant cells respond to deciliation with protein synthesis, another possible role of the Arp2/3 complex in ciliary assembly is the incorporation of a pool of existing proteins and membrane, which is actin-dependent (Jack et al., 2019). Further, disruption of Arp2/3 complex-mediated actin networks results in slow initial ciliary assembly, when it is likely that existing protein is being incorporated. We tested this by treating cells with cycloheximide (**Figure 2A, S2**) (Rosenbaum et al., 1969). Without protein synthesis, there is no trafficking or incorporation of new proteins. So, any ciliary growth we see is due to incorporation of existing protein alone. Normally, cells that are deciliated and treated with cycloheximide grow cilia to about half-length within 2 hours (**Figure 2B**). In *arpc4* mutant cells treated with cycloheximide, cilia display minimal growth; throughout a 5-hour period, only 6% of cells grew cilia (**Figure 2B**). This suggests the Arp2/3 complex is indispensable for incorporation of existing protein and membrane during ciliary assembly.

**Figure 2.**
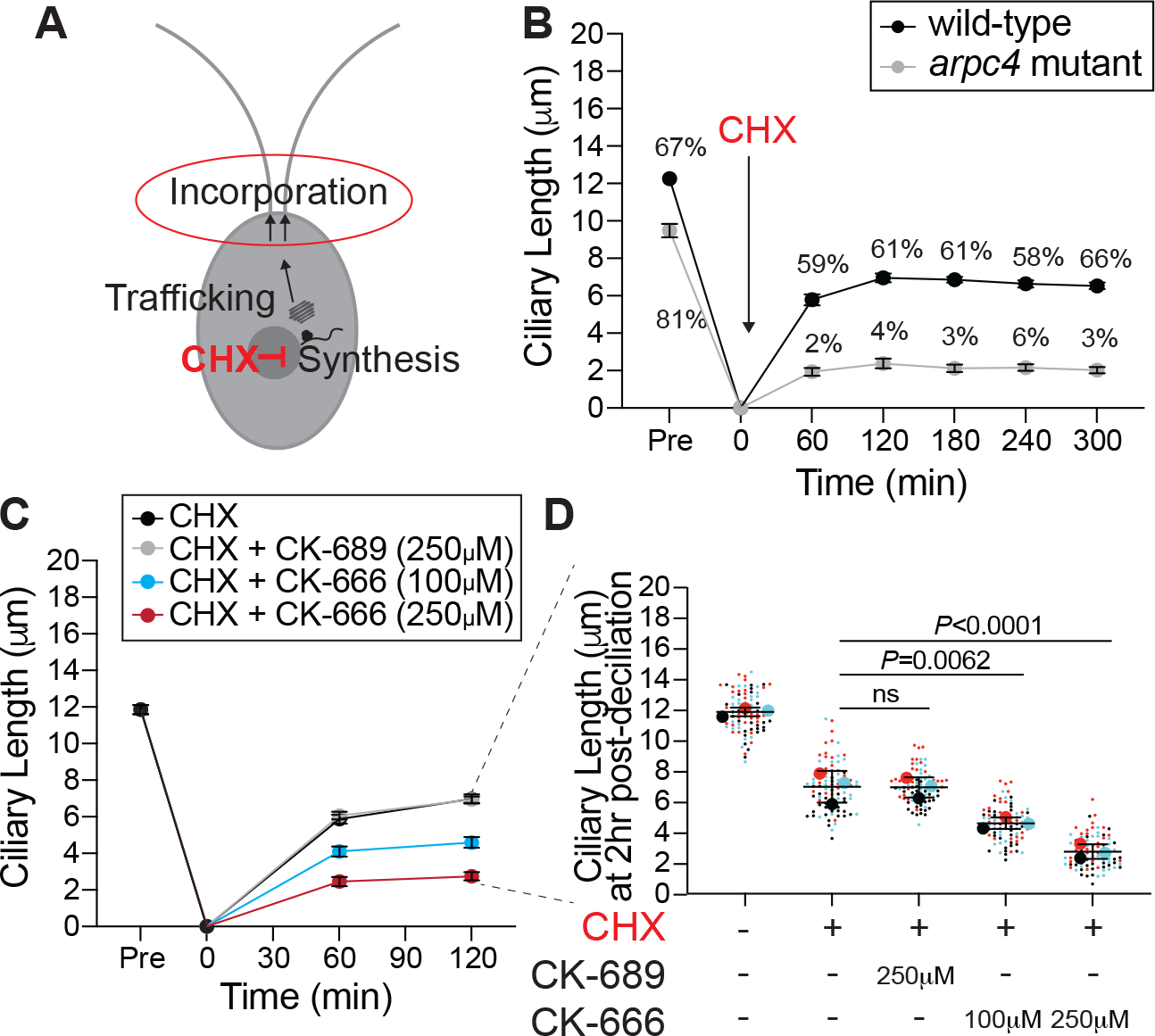
The Arp2/3 complex is required for incorporation of existing protein during ciliary assembly. **A)** Treating cells with cycloheximide inhibits protein synthesis, which means only incorporation of existing protein into the cilia is observed. **B)** Wild-type cells and *arpc4* mutants were deciliated and allowed to regrow in 10μM CHX. The percentages above the lines represent the percent of cells with cilia at the indicated time points. The mean is shown with error bars representing 95% confidence interval. n=30 for each strain and each time point in 3 separate experiments. For every time point besides 0 min, P<0.0001 for both length and percent ciliation. **C)** Wild-type cells were deciliated and then treated with a combination of 10μM cycloheximide (CHX) and CK-666 (100μM or 250μM) or CK-689 (the inactive control, 250μM) at the same concentration during regrowth. The mean is shown with error bars representing 95% confidence interval. n=30 for each strain and each time point in 3 separate experiments. At both 1 and 2 hour time points P<0.0001 for cells treated with CK-666 compared to wild-type cells, and ns for cells treated with CK-689 compared to wild-type cells. **D)** The length of cilia after 2 hours of treatment and regrowth. Superplot shows the mean of 3 separate experiments with error bars representing standard deviation. n=30 for each treatment group 3 separate experiments.

We suspected *arpc4* mutant cells either lacked the normal pool of ciliary precursor proteins or were unable to incorporate it. However, the inability of the genetic mutants to regenerate in cycloheximide prevents us from doing the typical studies testing new protein synthesis, precursor pool size, and new protein incorporation outlined in Jack et al. 2018 as they require regeneration in cycloheximide. To get around this, we used an acute perturbation, chemical inhibition in wild-type cells that have a normal ciliary precursor pool (as evidenced by their ability to grow to half-length in cycloheximide). Cells were deciliated and then CK-666 was added (in addition to cycloheximide) only for regrowth. Thus, the CK-666 could not affect precursor pool size. Cells treated with CK-666 and cycloheximide could not incorporate the precursor pool we know exists in these wild-type cells into cilia, while cilia of cells treated with only cycloheximide or cycloheximide and the inactive control, CK-689 grew to half length (**Figures 2C-D, S2**). This suggests the problem with incorporation in cells lacking a functional Arp2/3 complex lies outside of availability of the precursor pool.

### Cilia of *arpc4* mutant cells resorb faster in the absence of the Golgi

Because we see defects in ciliary assembly and maintenance when cells are likely incorporating existing protein, and we know the protein needed for assembly is in excess due to our acute perturbations with CK-666, we next investigated the membrane required for assembly. This is of particular interest as the Arp2/3 complex is often involved in membrane remodeling. Typically, the Golgi is thought to be the main source of membrane for cilia (Nachury et al., 2010; Rohatgi and Snell, 2010), and ciliary membrane, membrane proteins, and some axonemal proteins are transported in or attached to vesicles in cytosol (Wood and Rosenbaum, 2014). In *Chlamydomonas*, this has been demonstrated by the ciliary shortening of cells treated with Brefeldin A (BFA), a drug that causes Golgi collapse by interfering with ER to Golgi transport (Dentler, 2013). To determine if the Arp2/3 complex is involved in trafficking of new protein from the Golgi to cilia, we examined the Golgi following deciliation using transmission electron microscopy in *arpc4* mutants (**Figure S4A**). The Golgi appeared grossly normal, had the same number of cisternae, and did not show an accumulation of post-Golgi membrane as previously reported when perturbing filamentous actin (Jack et al., 2019) (**Figures S4A-B**).

Alternative pathways for delivery of ciliary material have also been found in *Chlamydomonas*. In one experiment, surface proteins were biotinylated and then cells were deciliated, so the membrane and proteins within cilia were lost. When cilia were allowed to regrow, biotinylated proteins were found within the new cilia suggesting they came from the plasma membrane (Dentler, 2013). Therefore, we hypothesized that due to its role in membrane remodeling in other organisms, the Arp2/3 complex may be part of an endocytic pathway that provides membrane to cilia (**Figure 3A**). To test if membrane could be coming from an endocytic source, we treated cells with BFA to collapse the Golgi and block exocytosis forcing cells to utilize other sources of ciliary membrane. Wild-type cilia treated with BFA resorb, but *arpc4* mutant cells resorb faster (**Figures 3B and D, S2**), and the number of cells with cilia in *arpc4* mutant cells dramatically decreased with BFA treatment (**Figure 3C**). Meanwhile, cells treated with other known ciliary resorption-inducing drugs that do not specifically target Golgi traffic, 3-isobutyl-1-methylxanthine (IBMX) (Pasquale and Goodenough, 1987) or sodium pyrophosphate (NaPPi) (Lefebvre et al., 1978) show an increased velocity of resorption in wild-type cells compared to *arpc4* mutant cells (**Figure S5**), so the faster resorption of *arpc4* mutant cells in BFA is specific to the effects of BFA. Thus, wild-type cells are more capable of maintaining cilia without membrane supply from the Golgi, suggesting there must be another source for membrane that is dependent upon the Arp2/3 complex, perhaps the cell body plasma membrane.

**Figure 3.**
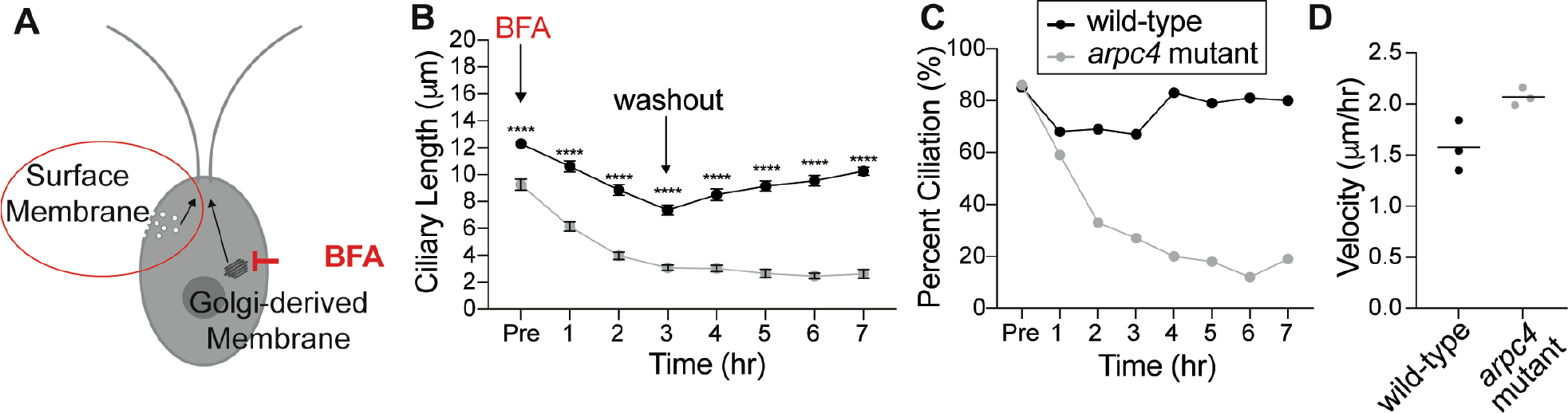
The Arp2/3 complex is required for ciliary maintenance in the absence of intact Golgi. **A)** Treating cells with Brefeldin A (BFA) causes the Golgi to collapse meaning any membranes and proteins used to maintain the cilia must come from other sources. **B)** Cells were treated with 36μM BFA for 3 hours at which time the drug was washed out. Wild-type is represented by black, while *arpc4* mutants are grey. The mean is shown with error bars representing 95% confidence interval. Error bars represent 95% confidence interval of the mean. n=30 for each time point and each strain in 3 separate experiments. **** represents P<0.0001. **C)** Percent ciliation of the cells in B. n=100. **D)** Resorption speed for wild-type cells and *arpc4* mutant cells as determined by fitting a line to the first 4 time points before washout and determining the slope of the line. Line represents the mean of 3 separate experiments. N=3. P=0.0314

### Ciliary membrane proteins follow different paths from the plasma membrane to the cilia

To determine if ciliary membrane and therefore membrane proteins could be coming from a pool in the plasma membrane we did an experiment first described in W. Dentler 2013. Surface proteins were biotinylated, then cells were deciliated. After the cilia regrew for 5 hours, they were isolated and probed for biotinylated protein (**Figure 4A**). Any biotinylated protein present in newly grown and isolated cilia must have come from a pool in the plasma membrane. First, we noticed differences in the biotinylated proteins found in wild-type cilia and *arpc4* mutant cilia before deciliation, suggesting there are overall differences in the composition of wild-type and *arpc4* mutant cilia (**Figure 4C**). If ciliary membrane proteins are coming from the cell body plasma membrane in an Arp2/3 complex-dependent manner as we hypothesize, this must be the case, as this would mean wild-type and *arpc4* mutant cells have differences in the trafficking pathways that bring ciliary material to cilia. More specifically, there are biotinylated proteins present in wild-type cells that are never present in *arpc4* mutant cells, so there is a mechanism for delivery of proteins to the cilia from the plasma membrane that absolutely requires the Arp2/3 complex (**Figures 4B-C**). There are also some proteins that are present to a higher degree in our *arpc4* mutants. We suspect this could be due to compensation by other pathways or defects in turnover of proteins in the cilia, perhaps through an exocytic mechanism. Next, looking at the cilia post-deciliation. Cilia were harvested 5 hours following deciliation because the *arpc4* mutant cilia regrow quite slowly. This means that cells might have time to employee other trafficking methods for getting material to cilia, but we still see striking differences. We found that while some proteins returned in both wild-type and *arpc4* mutant cells, some appeared to a lesser degree in *arpc4* mutant cells compared to wild-type cells (**Figures 8B-E, black arrow and black bars**) and some returned to a higher degree in *arpc4* mutant cells (**Figures 8B-E, grey arrow and grey bars**). This suggests there are multiple paths to the ciliary membrane, some of which are Arp2/3 complex-independent and some that are Arp2/3 complex-dependent. This may represent lateral diffusion and endocytosis respectively. Importantly, this assay tells us that membrane proteins can and do come to the cilia from the cell body plasma membrane.

**Figure 4.**
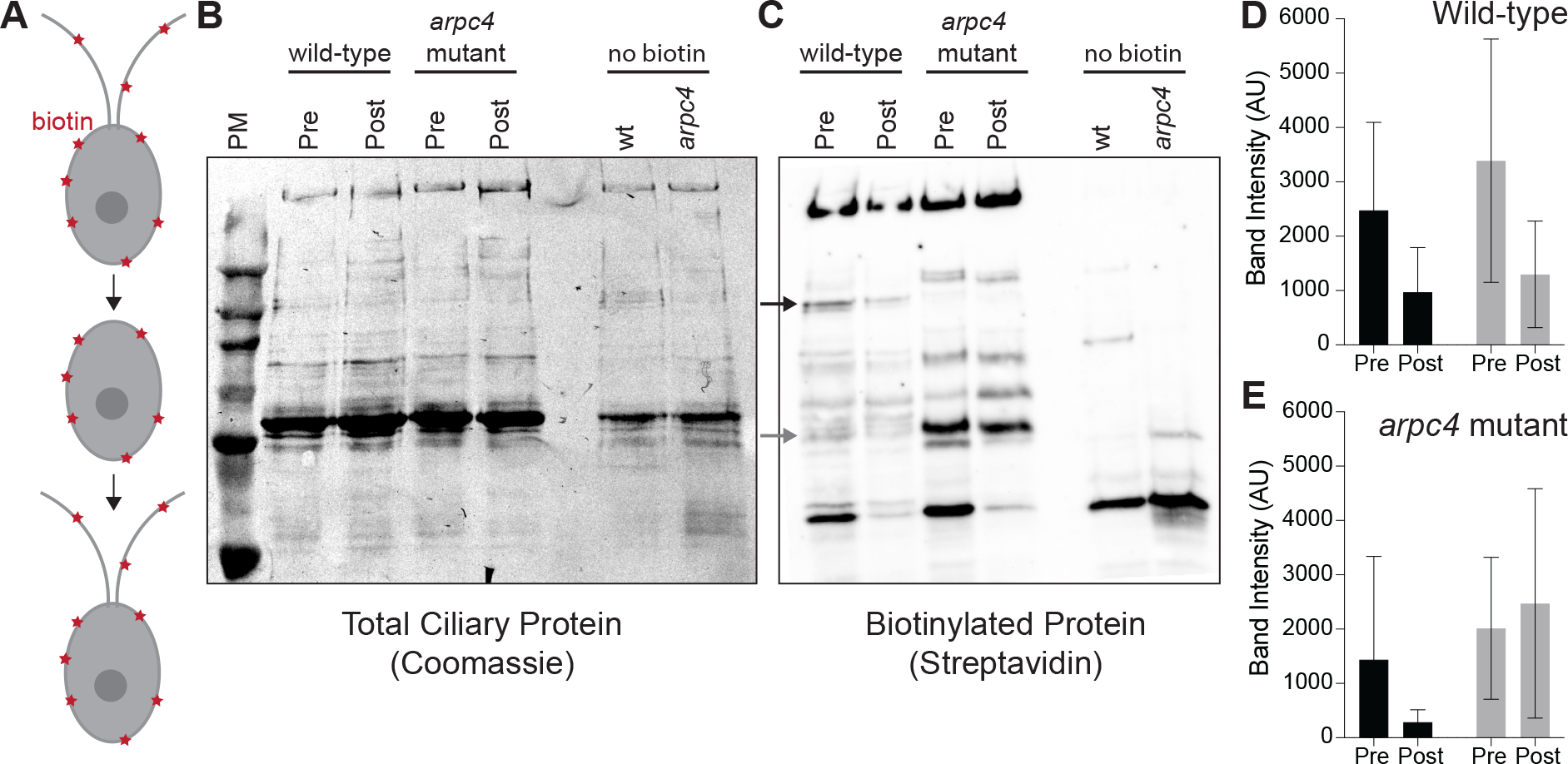
Ciliary membrane proteins have multiple paths from the plasma membrane. **A)** Cells were biotinylated, deciliated, and allowed to regrow before cilia were isolated and probed for biotinylated protein. **B)** Total protein in wild-type and *arpc4* mutant ciliary isolate investigated by western blot and Coomassie. **C)** Wild-type and *arpc4* mutant cells ciliary isolate was investigated by western blot and probed using streptavidin. Black arrow shows ciliary protein present to a higher degree in wild-type cells than *arpc4* mutant cells. Grey arrows show ciliary protein that is present to a higher degree in *arpc4* mutant cells than in wild-type cells. **D)** Bands represented by black and grey arrows are quantified for the wild-type cells. Data acquired from 3 separate experiments. **E)** Bands represented by black and grey arrows are quantified for the *arpc4* mutant cells. Data represented as the mean from 3 separate experiments. Error bars represent standard deviation.

### The Arp2/3 complex is required for the internalization and relocalization of a membrane protein from the periphery of the cell to cilia

Upon finding that ciliary membrane proteins can come from the cell body plasma membrane, we next asked if the internalization and relocalization of a known ciliary protein could be Arp2/3 complex dependent. SAG1 is a membrane protein important for mating in *Chlamydomonas* cells (Belzile et al., 2013; Ranjan et al., 2019). When cells are induced for mating with dibutyryl-cAMP (db-cAMP), SAG1 relocalizes from the cell periphery to cilia, where it facilitates ciliary adhesion between mating cells. This relocalizaiton of SAG1 is thought to occur through internalization and internal trafficking on microtubules (Belzile et al., 2013; Ranjan et al., 2019).

We examined whether actin and the Arp2/3 complex were required for transport of HA-tagged SAG1 to the cell apex and cilia during mating (**Figure 5A**). We observed cells treated with either LatB to depolymerize IDA5 or CK-666 to perturb the Arp2/3 complex (**Figures 5, S2**). Before induction, SAG1-HA localized to the cell periphery (**Figure 5B, top**). 30 minutes after induction with db-cAMP, SAG1-HA relocalized to the cell apex and to cilia in untreated cells (**Figure 5B, left**). In both LatB and CK-666 treated cells, this apical enrichment decreased (**Figure 5B, middle and right**). We took line scans through the cell from the apex to the basal region (**Figures 5C-D**) and calculated the percentage of cells with apical enrichment. Untreated cells had a higher percent of apical enrichment when compared with LatB or CK-666 treated cells (**Figure 5E**). Thus, cells with perturbed Arp2/3 complex or filamentous actin show decreased efficiency of SAG1-HA relocalization.

**Figure 5.**
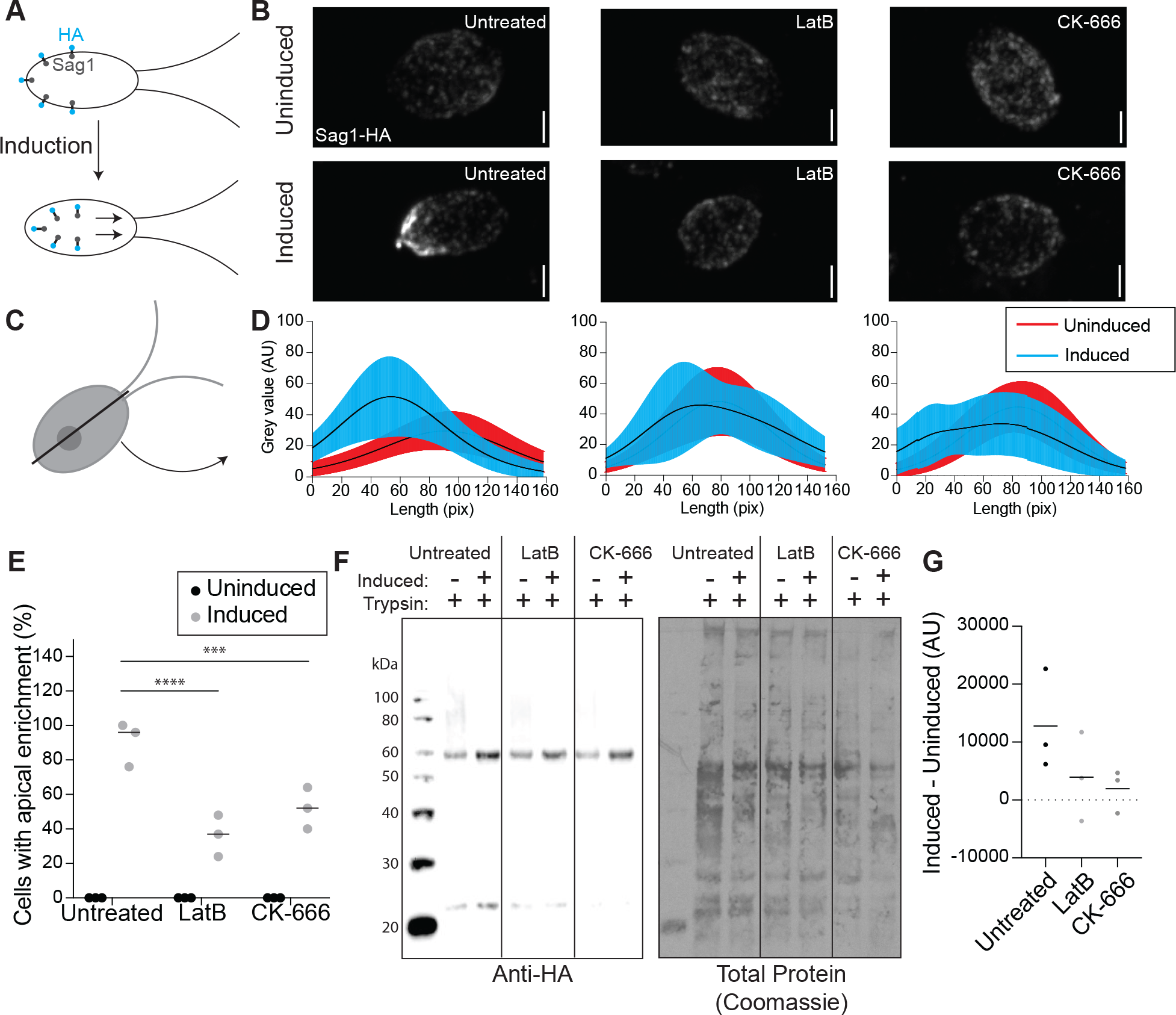
The Arp2/3 complex is required for the relocalization and internalization of the ciliary protein SAG1 for mating. **A)** When mating is induced SAG1-HA is internalized and relocalized to the apex of the cells and cilia for agglutination. **B)** Maximum intensity projections of z-stacks showing SAG1-HA. Scale bar represents 2μm. **C)** Line scans were taken through the cells in z-stack sum images. **D)** Line scans in untreated cells (left), LatB treated cells (middle), and CK-666 (right) were normalized and fit with a gaussian curve. The curves were averaged. Black lines represent mean and shaded regions represent standard deviation. Red represents uninduced samples, cyan represents induced samples. 0 on the y-axis represents the apical region of the cell. n=30 from a single representative experiment. **E)** Percentage of cells with apical enrichment for uninduced (black) and induced (grey) cells for each treatment group. The mean is shown with the solid line. N=30 for 3 separate experiments for each treatment. **F)** Western blot showing amount of SAG1-HA in uninduced and induced cells in each treatment group all treated with 0.01% trypsin. **G)** Intensity of the bands in F were normalized to the total protein as determined by Coomassie staining and quantified in ImageJ was used to subtract uninduced from induced to give a representation of the amount of SAG1-HA internalized with induction. Line represents mean of 3 separate experiments.

We asked if this decrease in relocalization in cells with actin and Arp2/3 complex inhibition could be due to a decrease in internalization of SAG1-HA through a process that seems to require endocytosis. We used a method first described by Belzile et al. 2013, where cells were induced and treated with a low percentage (0.01%) of trypsin, which hydrolyzes exterior proteins but cannot enter the cell. In untreated cells, we see an increase in SAG1-HA protein levels following induction because SAG1-HA is internalized and becomes protected from trypsin (**Figure 5F**). In cells treated with either LatB or CK-666 we see a decrease in this trypsin protection (**Figure 5F**). We quantified this by subtracting the amount of protein before induction from the amount of protein present after induction, which gives a value representing the amount of SAG1-HA protected from trypsin due to internalization (**Figure 5G**). The decrease in SAG1-HA following induction in LatB or CK-666 treated cells indicates a role for Arp2/3 complex and actin in internalization of this specific ciliary membrane protein.

### Apical actin dots are dependent on the Arp2/3 complex

Since ciliary membrane proteins can come from the Golgi or the plasma membrane and *arpc4* mutant cells have a more severe defect in incorporating ciliary proteins from non-Golgi sources, we asked if Arp2/3 complex-mediated actin networks might be responsible for plasma membrane remodeling in *Chlamydomonas* as it is in other organisms. Thus, we looked at the effects of loss of Arp2/3 complex function on actin structures. Using new protocols for visualizing actin in *Chlamydomonas* (Craig et al., 2019), we stained wild-type cells and *arpc4* mutant cells with fluorescent phalloidin. In wild-type cells, apical dots reminiscent of endocytic actin patches in yeast are seen near the base of cilia (**Figure 6A**). We quantified the presence of dots in the wild-type cells compared to *arpc4* mutant cells (**Figure 6A-B**). While about 70% of wild-type cells contain the dots, less than 5% of *arpc4* mutant cells had dots (**Figure 6B**), suggesting the Arp2/3 complex is required for formation of this structure. Expression of the ARPC4-V5 construct in *arpc4* mutant cells rescued the dots (**Figure 6C**). Because ARPC4-V5 staining showed diffuse signal throughout the cell, we are not able to determine whether or not active Arp2/3 complex localizes to the dots (**Figure S1C**). However, the reliance of this structure on the Arp2/3 complex suggests that the Arp2/3 complex is definitely involved in this structure. This led us to question whether these dots could represent membrane remodeling.

**Figure 6.**
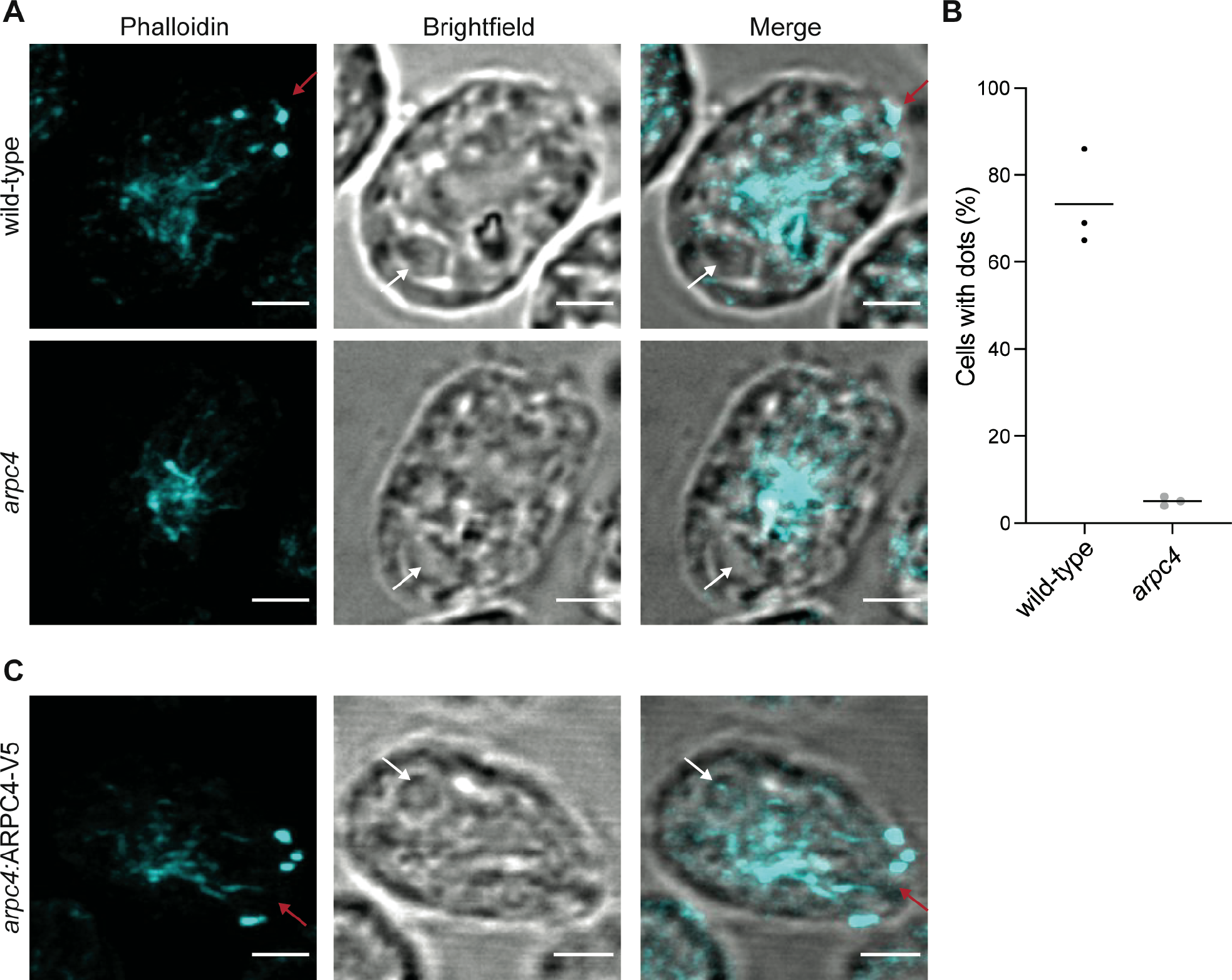
Loss of a functional Arp2/3 complex results in changes in actin distribution. **A)** Wild-type and *arpc4* mutant cells stained with phalloidin to visualize the actin network along with brightfield to show cell orientation. Images were taken as a z-stack using airsycan imaging and are shown as a maximum intensity projection. Red arrow is pointing to dots at the apex of the cell, and white arrow is pointing to the pyrenoid near the basal end of the cell. Scale bars represent 2μm. **B)** Percentage of cells with apical dots as shown in A. Percentages taken from 3 separate experiments where n=100. Line represents the mean. P<0.0001. **C)** Presence of apical dots in the *arpc4* mutant rescue expressing ARPC4-V5. Images were taken as a z-stack using airsycan imaging and are shown as a maximum intensity projection. Red arrow is pointing to dots at the apex of the cell, and white arrow is pointing to the pyrenoid near the basal end of the cell. Scale bars represent 2μm.

### Endocytosis occurs in *Chlamydomonas*

The Arp2/3 complex is thought to be involved in endocytosis in cell-walled yeast to overcome turgor pressure (Aghamohammadzadeh and Ayscough, 2009; Basu et al., 2014; Carlsson and Bayly, 2014). *Chlamydomonas* cells also have a cell wall and since the actin dots resemble yeast endocytic pits (Adams and Pringle, 1984; Ayscough et al., 1997; Goode et al., 2015), we hypothesized that Arp2/3 complex-dependent endocytosis might be occurring in *Chlamydomonas* though this process has not yet been directly demonstrated in this organism. To determine what kind of endocytosis likely occurs in these cells, we compared the endocytosis-related proteins found in mammals and plants to those in *Chlamydomonas* (**Figure 7A**). *Chlamydomonas* lacks much of the important machinery for almost all typical endocytosis processes, including caveolin for caveolin-mediated endocytosis, flotillin for flotillin-dependent endocytosis, and endophilin for endophilin-dependent endocytosis (**Figure 7A**). However, clathrin-mediated endocytosis is conserved to a higher extent than other endocytic mechanisms.

**Figure 7.**
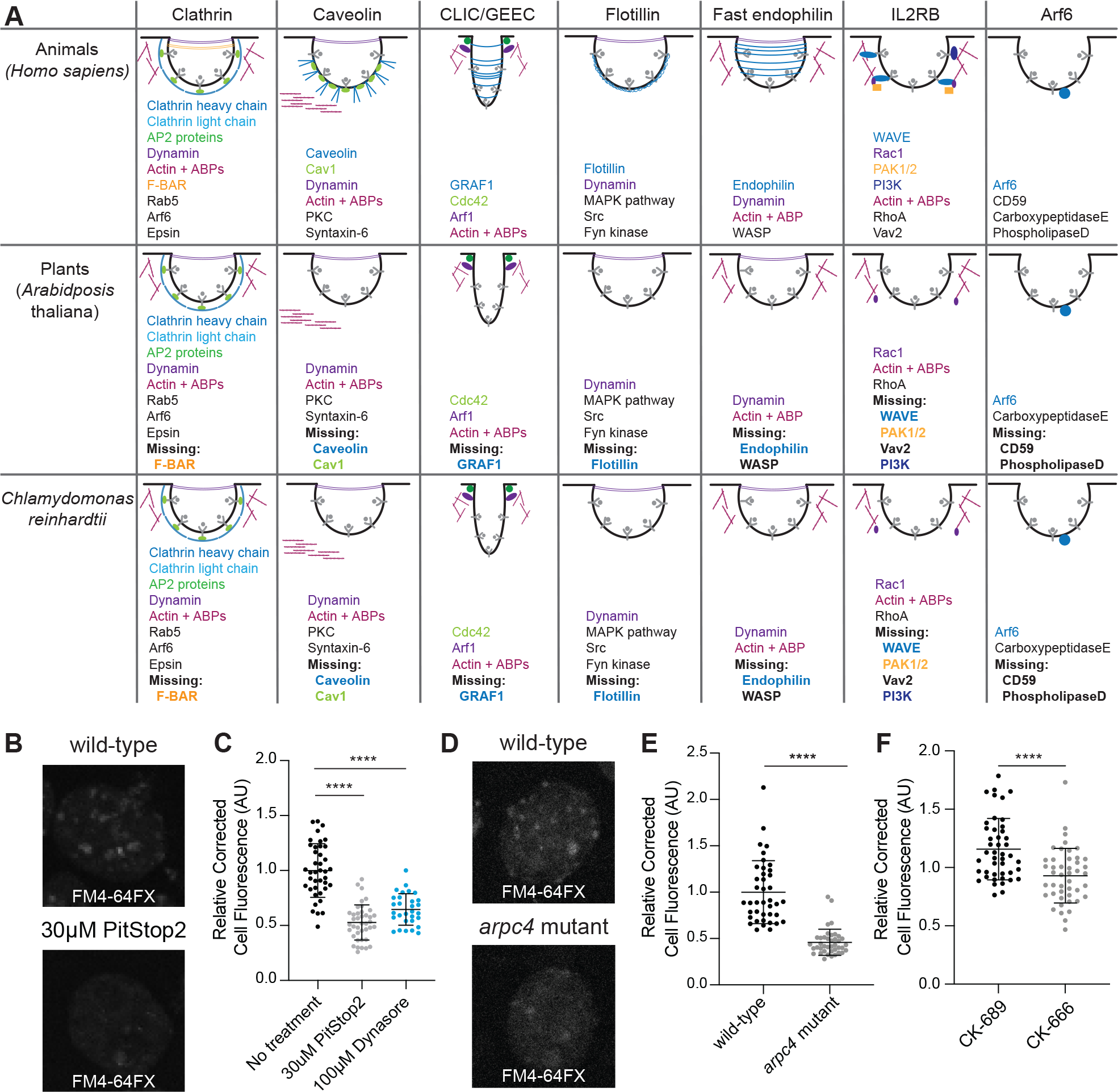
Arp2/3 complex-dependent endocytosis is conserved in *Chlamydomonas*. **A)** Gene presence was determined using BLAST. Word colors correspond to diagram colors. **B)** Cells treated with 30μM PitStop2 were incubated with FM4-64FX and imaged on a spinning disk confocal. Max intensity projections of z-stacks are shown. Scale bars are 2μm. **C)** The background corrected fluorescence for each sample, including cells treated with 100μM Dynasore. The mean is shown with error bars showing standard deviation. n=30 in 3 separate experiments. P<0.0001. **D)** Wild-type and *arpc4* mutant cells treated with FM4-64FX and imaged on a spinning disk confocal. Max intensity projections of z-stacks are shown. Scale bars are 2μm. **E)** The background corrected fluorescence for each sample. The mean is shown with error bars representing standard deviation. n=30 in 3 separate experiments. P<0.0001. **F)** The background corrected fluorescence for cells treated with CK-666 or CK-689. The mean is shown with error bars representing standard deviation. n=30 in 3 separate experiments. P<0.0001.

We aimed to probe the likelihood of endocytosis occurring in *Chlamydomonas*, but a mutant for the proteins involved in clathrin-mediated endocytosis does not currently exist and methods of targeted mutagenesis in *Chlamydomonas* are not yet reliable. So, we turned to our best alternative PitStop2, which inhibits the interaction of adaptor proteins with clathrin, halting clathrin endocytosis, despite reported off-target effects on global endocytosis in mammalian cells (Willox et al., 2014) (**Figure S2**). We also used the dynamin inhibitor Dynasore, which is thought to block endocytosis by inhibiting the GTPase activity of dynamin (Macia et al., 2006), although this inhibitor has also been found to affect actin in some mammalian cells (Mooren Olivia L. and Schafer Dorothy A., 2009; Park et al., 2013; Yamada et al., 2009). Although both PitStop2 and Dynasore are reported to have off-target effects in different pathways, their intended target is in the same pathway. Therefore, by using both we hope to reduce concerns of off-target effects. To further minimize off-target effects, this experiment was done in a fast time scale and at the lowest concentration possible. For this experiment, we used the fixable lipophilic dye FM 4-64FX (Cochilla et al., 1999; Gachet and Hyams, 2005), which is impermeable to the plasma membrane but is endocytosed into cells showing bright foci where dye is enriched in endocytic compartments. We incubated the dye for 1 minute to allow enough time for internalization into endosomes but not enough for incorporation into various cellular membrane structures. The ability of cells to internalize membrane was measured by calculating the total cell fluorescence inside the cell after dye internalization (**Figure 7B**). Cells treated with PitStop2 or Dynasore internalized significantly less membrane dye (**Figure 7C**), which supports the idea that endocytosis is occurring in these cells and that it is likely clathrin-mediated.

Next, we tested whether endocytosis is Arp2/3 complex-dependent by using this membrane internalization assay on *arpc4* mutant cells compared to wild-type cells and CK-666 treated cells compared to CK-689 treated cells. *arpc4* mutant cells and cells treated with CK-666 have decreased total cell fluorescence (**Figure 7D-F**) suggesting endocytosis in *Chlamydomonas* is Arp2/3 complex-dependent.

### Actin dots increase in an Arp2/3 complex and endocytosis-dependent manner following deciliation

Having established that the Arp2/3 complex is required for ciliary assembly, membrane dye internalization, and endocytosis of a known ciliary protein, we wondered if these functions could be connected given that *arpc4* mutant cells have defects in maintaining cilia from non-Golgi sources. We returned to the Arp2/3 complex-dependent actin dots that are reminiscent of endocytic pits in yeast. Because ciliary membrane and proteins can come from the plasma membrane (Dentler, 2013), we suspected there would be an increase in actin dots following deciliation. We used phalloidin to visualize the actin cytoskeleton of wild-type cells before and immediately following deciliation, as well as 10 minutes later (**Figure 8A**). We saw an increase in both the percentage of cells with dots and the number of dots per cell immediately following deciliation that returned to normal by 10 minutes (**Figures 8A, D**). This is consistent with the results in **Figures 1E-F** and confirms that the defect in ciliary assembly is due to an event occurring very early in ciliary assembly, within the first few minutes after deciliation.

In the *arpc4* mutant cells dots were never observed, before or after deciliation (**Figure 8B, D**), confirming these dots are Arp2/3 complex dependent. Next, we investigated if the dots were due to endocytosis by treating cells with PitStop2 and looking for this same increase in dots. This treatment almost fully blocked the appearance of dots following deciliation and eliminated the presence of cells with 3 or more dots (**Figure 8C-D**), suggesting an Arp2/3 complex-dependent endocytic mechanism is related to these dots that occur immediately following endocytosis when ciliary material is in high demand.

## DISCUSSION

In this study, we investigate the Arp2/3 complex of *Chlamydomonas reinhardtii* that functions to maintain and assemble cilia. This complex potentially lacks the ARPC5 subunit, although it is possible that a divergent ARPC5 exists. In yeast, deletion of any of the genes encoding Arp2/3 complex members causes severe defects, but these defects differ in severity depending on the complex members deleted, suggesting complex members have varying degrees of importance in Arp2/3 complex function (Winter et al., 1999). The role of ARPC5 in actin nucleation is being investigated, but some groups have found it unnecessary for function of the complex (Gournier et al., 2001; von Loeffelholz et al., 2020). Furthermore, our data show that knocking out function of the ARPC5-less *Chlamydomonas* Arp2/3 complex results in various phenotypes, suggesting the wild-type complex is active. In this paper, we study the Arp2/3 complex using 2 main perturbations: genetic inhibition of the ARPC4 member of the complex and chemical inhibition with the inhibitor CK-666. CK-666 is designed for mammalian cells, but we believe that CK-666 is functional in *Chlamydomonas* because it can recapitulate the effects of the genetic mutant. Regardless, we used both the genetic perturbation and the chemical perturbation for nearly every experiment looking at the role of the Arp2/3 complex in these phenotypes. Additionally, if we treat the *arpc4* mutant with CK-666 we do not see these same phenotypes, again suggesting that CK-666 is acting to block the same functions as the genetic arcp4 mutant (**Figure 1A**). Because the Arp2/3 complex has known functions in membrane dynamics and because of our data demonstrate a role for the Arp2/3 complex of *Chlamydomonas* in membrane and membrane protein internalization, this led us to pursue models of Arp2/3 complex-dependent membrane trafficking to cilia.

Previously, three models of membrane protein trafficking to cilia have been proposed regarding where ciliary vesicles fuse relative to a diffusion barrier composed of septins (Hu Qicong et al., 2010), which delineates ciliary membrane and cell body plasma membrane (Nachury et al., 2010). The first is that Golgi vesicles containing ciliary proteins fuse with the ciliary membrane inside the cilium. Proteins, both membrane and soluble, have been found to travel from the Golgi to the cilia on or in cytoplasmic vesicles (Wood and Rosenbaum, 2014). Second, Golgi vesicles containing ciliary proteins fuse outside but near the cilium still within the diffusion barrier (Nachury et al., 2007; Papermaster et al., 1985; Zuo et al., 2009). In *Chlamydomonas*, mastigoneme proteins travel from the Golgi and are exocytosed for use on the exterior of the cell (Bouck, 1971). In the third model, Golgi vesicles containing proteins fuse with the plasma membrane outside the diffusion barrier where they move in the plane of the plasma membrane across this barrier, perhaps through lateral diffusion that requires remodeling or passing through the diffusion barrier. Evidence for this path was shown using Hedgehog signaling protein Smoothened, which relocalizes in a dynamin-independent manner from the plasma membrane to the cilia immediately after stimulation in pulse labeling studies (Milenkovic et al., 2009).

Our data all together support a fourth model, likely occurring in concert with other models, in which membrane and membrane proteins are recruited to the cilium from a reservoir in the cell body plasma membrane. We show that the Arp2/3 complex is required for ciliary assembly from zero-length (**Figure 1**); we show that ciliary membrane proteins can and do come from the cell body plasma membrane, both generally (**Figure 4**) and for a specific protein (**Figure 5**); we show that the Arp2/3 complex is required for endocytosis (**Figure 7**) and for the formation of actin dots reminiscent of endocytic pits or patches (**Figures 6, 8**); and finally we show an endocytosis-dependent increase in Arp2/3 complex-mediated actin dots immediately following deciliation (**Figure 8**). Thus, we hypothesize that ciliary membrane proteins and membrane targeted to the plasma membrane of the cell outside the diffusion barrier can be endocytosed and trafficked to cilia, either within or outside of the diffusion barrier in an actin and Arp2/3 complex-dependent manner.

Although our data do not eliminate the possibility of Arp2/3 complex function in supply of ciliary membrane and protein stored in other endosomal compartments, ciliary localization of proteins initially labeled on the cell surface with biotin (**Figure 8**) suggests some ciliary membrane proteins incorporated during assembly are coming from the plasma membrane itself. One limitation to this study is the time frame. We isolated cilia following 5 hours of regrowth to get a measurable amount of *arpc4* mutant cilia, which have very defective growth. This means that compensatory mechanisms such as synthesis and slower trafficking may be involved. We also cannot rule out an additional role for the Arp2/3 complex in delivery of existing soluble proteins to cilia. An in-depth analysis of soluble protein recruitment and incorporation would be a useful next step to determine if the Arp2/3 complex is involved in other ciliary assembly related processes. Our data also does not preclude Arp2/3 complex function in other membrane dynamics. However, all the data in this paper together support a model involving membrane remodeling and endocytosis. An endocytic mechanism of trafficking in intracellular ciliogenesis has been investigated in mammalian RPE1 cells. The ciliary pocket found at the base of primary and motile cilia formed intracellularly has been found to be an endocytically active region (Molla-Herman et al., 2010) but clathrin-mediated endocytosis was not required for ciliogenesis in those cells. The Bardet Biedl Syndrome complex (BBsome), which is involved in regulation of ciliary membrane protein composition, has been shown to interact with clathrin directly at the ciliary pocket to facilitate membrane sorting in trypanosomes (Langousis et al., 2016). Further, some BBsome complex members resemble coat proteins such as clathrin (Jin et al., 2010) suggesting a direct role for the this cilium regulatory complex in membrane functions. It has also been found that disruption of recycling endosomes reduces the localization of polycystin-2 to cilia, suggesting a role for recycling endosomes in the localization of proteins to the cilia (Monis et al., 2017). In *Chlamydomonas*, clathrin heavy chain has been found to localize at the base of cilia (Kaplan et al., 2012). While the mechanism was unknown, it has been shown that plasma membrane surface-exposed proteins are relocalized to cilia during ciliary regeneration (Dentler, 2013), a result we recapitulated and demonstrated depends, in part, upon the Arp2/3 complex.

**Figure 8.**
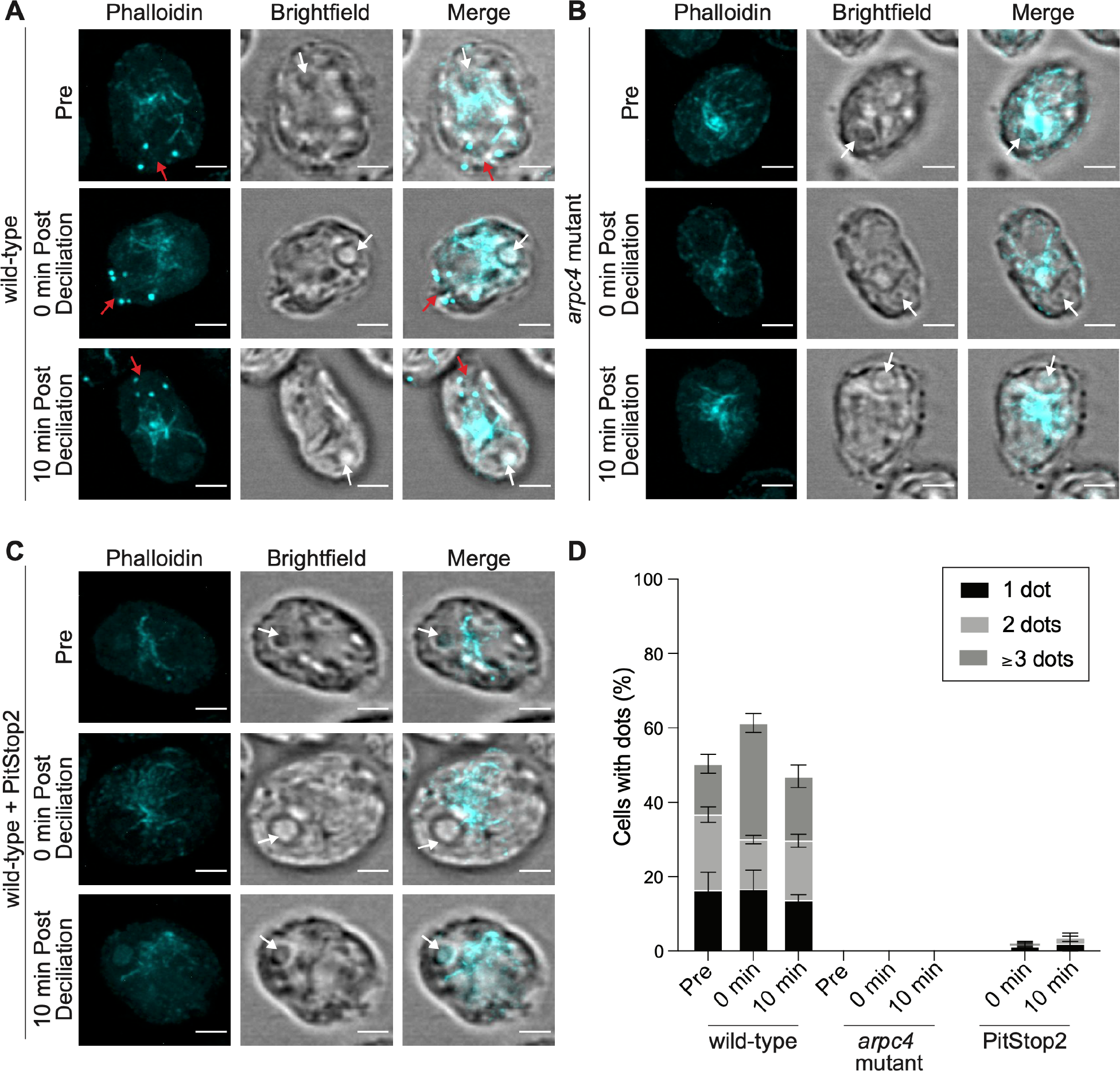
Actin dots require the Arp2/3 complex and endocytosis. **A-C)** Wild-type cells **(A)**, *arpc4* mutant cells **(B)**, and wild-type cells treated with 30 μM PitStop2 **(C)** stained with phalloidin to visualize the actin network before deciliation, immediately following deciliation, and 10 minutes following deciliation. Brightfield images are to visualize cell orientation. Images were taken as a z-stack and are shown as a maximum intensity projections. Scale bar represents 2μm. Red arrows point to dots at the apex of the cell, and white arrows point to the pyrenoid at the opposite end of the cell. **D)** The percentage of cells with 1 dot, 2 dot, or 3 dots in each condition. Quantification based on sum slices of z-stacks taken using a spinning disk confocal. n=100 in 3 separate experiments. For wild-type, the total number of cells with dots is significantly different for the 0 min time point (**) and the number of dotted cells with 3 or more dots is significantly different for the 0 time point (****).

Altogether, this leads us to hypothesize that the role of the Arp2/3 complex in ciliary assembly is through endocytic recruitment from a ciliary protein reservoir in the plasma membrane before newly synthesized protein and Golgi-derived membrane can supply additional materials (**Figure 9B**). While this model provides a possible route that some ciliary proteins and membranes take to the cilia, we believe this is one of several paths that can be taken to the cilia. This could be further investigated by determining specific proteins that may take these different paths to the cilia. Trafficking to cilia is likely cargo- and time-dependent, and which path proteins take may tell us the order and speed in which they populate the cilium for subsequent function.

**Figure 9.**
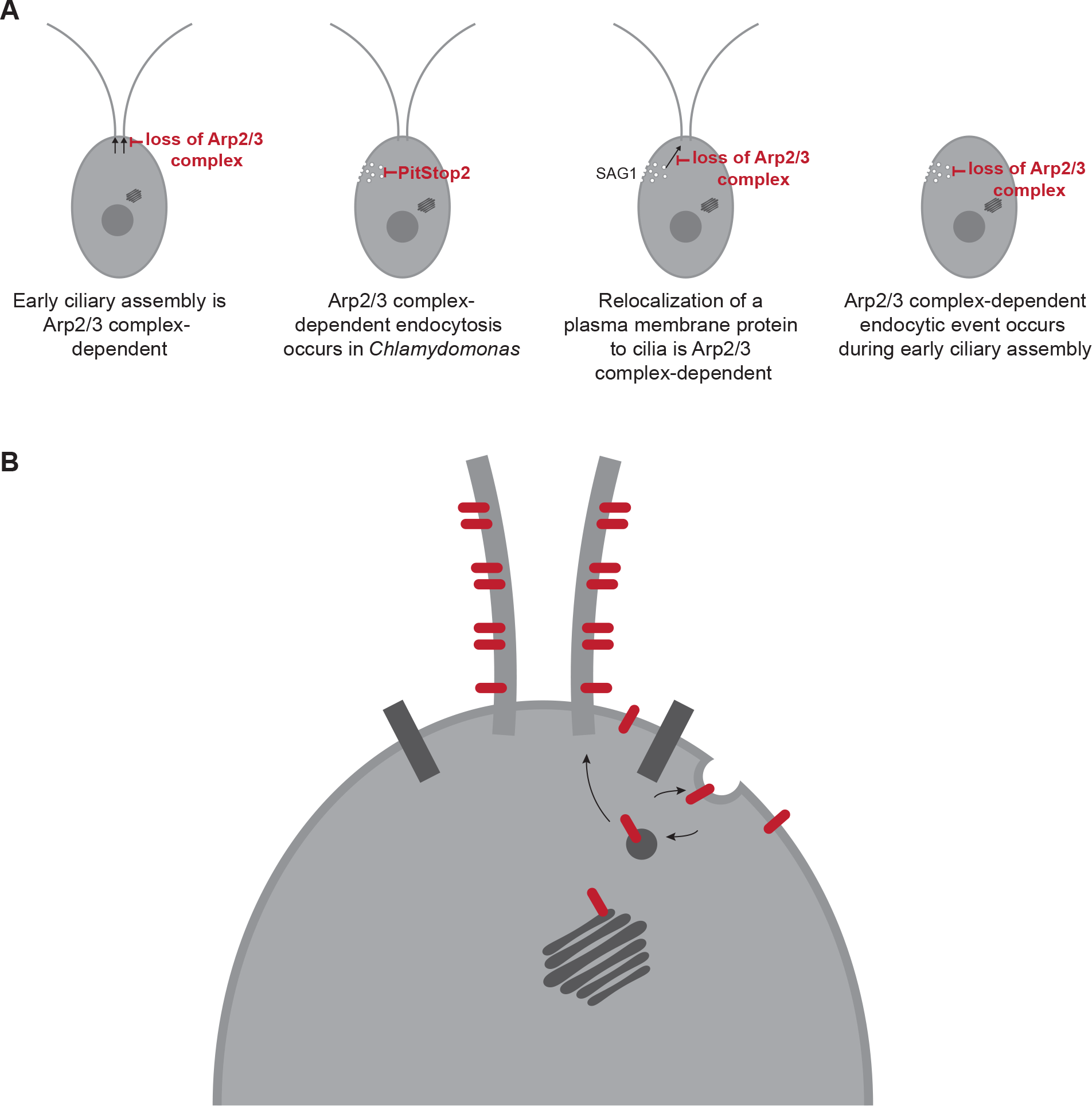
The Arp2/3 complex is required for membrane and protein delivery via a Golgi-independent, endocytosis-like process. **A)** Arp2/3-mediated actin networks are required for ciliary assembly in *Chlamydomonas* particularly during the initial stages. These actin networks are also required for endocytosis, and for the endocytosis-like relocalization of a ciliary protein from the plasma membrane to the cilia. Finally, a large endocytic event occurs immediately following deciliation that is Arp2/3 complex-mediated. **B)** Proposed model of membrane protein and membrane transport from the plasma membrane to the cilia through endocytosis.

## METHODS

### Strains

The wild-type *Chlamydomonas* strain (CC-5325) and the *arpc4* mutant (LMJ.RY0402.232713) are from the *Chlamydomonas* resource center. The *arpc4:*ARPC4-V5 strain was made by cloning the gene into pChlamy4 (*Chlamydomonas* resource center). Colonies were screened for the absence (in the case of the mutant) or presence (in the case of the rescue) by PCR using the primers AAAAGAATTCATGGCGCTCTCACTCAGGCCATA and AAAATCTAGACAGAAGGCAAGGGAGCGCAGGAA. The SAG1-HA strain was a gift from William Snell. Cells were grown and maintained on 1.5% Tris-Acetate Phosphate Agar (TAP) plates (*Chlamydomonas* resource center) under constant blue (450-475 nm) and red light (625-660 nm). For experiments, cells were grown in liquid TAP media (*Chlamydomonas* resource center) overnight under constant red and blue light with agitation from a rotator. To induce gametes for mating for the SAG1-HA experiments, cells were grown in liquid M-N media (*Chlamydomonas* resource center) overnight with constant red and blue light and agitation.

### Ciliary studies

For steady state experiments, cells were treated with specified drugs [either 100μM CK-666, 250μM CK-666 (Sigma, 182515) 250μM CK-689 (CalBiochem, 182517), 10μM LatB (Sigma, L5288), 10μM CHX (Sigma, C1988), or 36μM BFA (Sigma, B7651) all diluted in DMSO (Sigma, D2650)] and incubated with agitation for the allotted times. Following any incubation (as well as a pre sample), cells were diluted in an equal volume of 2% glutaraldehyde (EMS, 16220) and incubated at 4º Celsius until they sediment (within 24hrs). Following sedimentation, cells were imaged using a Zeiss Axioscope 5 DIC microscope with a 40X objective (0.75 numerical aperture) at room temperature with no immersion media or imaging media. Images were acquired using Zeiss Zen 3.1 (blue edition). Cilia were then measured using the segmented line function in ImageJ. One cilium per cell was measured and 30 cilia total were measured.

For regeneration experiments, a pre sample was taken by adding cells to an equal volume of 2% glutaraldehyde. Then cells were deciliated with 115μL of 0.5N acetic acid for 45 seconds. After this short incubation, the pH was returned to normal by adding 120μL of 0.5N KOH. A 0-minute sample was again taken by adding cells to an equal volume of 2% glutaraldehyde. Then cells were incubated with agitation and allowed to regrow cilia for the allotted time period with samples taken at the indicated time points by adding cells to an equal volume of 2% glutaraldehyde. Cells in glutaraldehyde were allowed to incubate at 4º Celsius until sedimentation (within 24hrs). Then, cells were imaged using the same Zeiss DIC microscope with a 40X objective and the same software. Cilia were then measured using the segmented line function in ImageJ. One cilium per cell was measured and 30 cilia total were measured.

### ARPC4 Rescue

The ARPC4 genetic sequence was isolated from Chlamydomonas cDNA using PCR with the Q5 DNA Polymerase (NEB, M0491L). The resulting fragment and the pChlamy4 plasmid (Thermofisher, A24231) were digested with EcoRI (NEB, R0101S) and XhoI (NEB, R0146S) for 1 hour followed by heat inactivation. Then, the fragment and vector were mixed in a 5:1 ratio and ligated with T4 DNA Ligase (NEB, M0202L) overnight at 16°C. The vector was then transformed into One Shot TOP10 chemically competent cells (Invitrogen, C404003) following the product protocol. The transformed competent cells were plated on LB plates with 100 μg/mL ampicillin (IBI Scientific, IB02040) and grown overnight at 37°C. The following morning colonies were screened using DreamTaq DNA Polymerase (Thermo, EP0702) and the forward primer AAAAGAATTCATGGCGCTCTCACTCAGGCCATA and the reverse primer AAAATCTAGACAGAAGGCAAGGGAGCGCAGGAA. Positive colonies were grown in liquid LB with 100 μg/mL ampicillin overnight at 37°C. Plasmid DNA was isolated from bacterial cells and sequenced.

Plasmids containing the ARPC4 DNA were then transformed into *Chlamydomonas* cells. First, a 5 mL liquid culture of *arpc4* mutant cells was grown overnight with agitation and constant light in TAP. The following day 25 mL of TAP was brought to an OD_730_ of 0.1 using the 5 mL culture. This was incubated with agitation and under constant light overnight. The culture reached an OD_730_ of 0.3-0.4 for the transformation. Once this occurred, the plasmid was linearized using ScaI (NEB, R3122L). Meanwhile, the 25 mL culture was centrifuged at 500xg to pellet the cells. The TAP was removed and replaced with 5 mL of Max Efficiency Transformation Reagent for Algae (Invitrogen, 100021485). This was repeated 2 times. After the final centrifugation, the cells were resuspended in 250 μL of Max Efficiency Transformation Reagent. This was then split in two. 1000 μg of linearized plasmid was added to each. This was then electroporated using a BioRad Electroporator at 500V, 50 μF, and 800 Ω in a 4 mm cuvette. The cells were removed from the cuvette following electroporation, suspended in 7 mL of TAP + 40 mM sucrose, and incubated overnight in the dark. The following day the cells were pelleted and streaked on TAP + Zeocin (10 μg/mL) plates, then incubated in constant light for approximately 1 week or until colonies formed.

Colonies were screened using DreamTaq DNA Polymerase (Thermo, EP0702) and the same primers as above. Positive colonies were streaked onto new plates and allowed to grow up. Expression of ARPC4-V5 was confirmed with a western blot. Liquid cultures of cells were grown overnight, then pelleted at 500 xg for 1 minute. Cells were resuspended in lysis buffer [5% glycerol (), 1% NP-40 (), 1mM DTT (), 1X protease inhibitors (Thermo, 1861281)] and lysed using bead beating. Cell debris was spun down at 14000xg for 15 minutes. An equal amount of protein was loaded to a NuPAGE 10% Bis-Tris gel (Invitrogen, NP0316). The resulting gel was transferred to PVDF membrane (Millipore, IPVH00010) which was then blocked with 5% milk in PBST. The blot was incubated with rabbit anti-V5 primary antibody (Cell Signaling, D3H8Q) diluted to 1:1000 in 1% BSA, 1% milk overnight at 4°C to probe for V5. The following day blots were washed 3 times in 1X PBST, then incubated with HRP-conjugated goat anti-rabbit secondary (Thermo, G-21234) diluted to 1:5000 in 1% milk 1% BSA for 1 hour at room temperature. The blot was washed again 3 times with 1X PBST. Then the blot was probed with West Pico Chemiluminescent Pico Substrate (Invitrogen, 34580). The same membrane was stripped of antibody and total protein was determined with Coomassie (Sigma, B0149) staining. Band intensity was measured in ImageJ and normalized to total protein.

### Click-iT OPP Protein Synthesis Assay (Invitrogen, C10457)

Cells were grown overnight in TAP. The following day cells were deciliated as described above and allowed to regrow either with or without cycloheximide (10μM) to block protein synthesis. 1 hour following deciliation, cells were mounted onto poly-lysine (EMS, 19321-B) coverslips. Cells on coverslips were incubated with Click-iT OPP reagent containing the O-propargyl-puromycin (OPP) which is incorporated into nascent polypeptides for 30 minutes. OPP was removed and cells were washed once in PBS. Cells were then fixed with 4% PFA (EMS, 15710) in 1X HEPES (Sigma, 391338) for 15 minutes, then permeabilized with 0.5% Triton-X 100 in PBS for 15 minutes. Cells were washed twice with PBS. Detection was performed by incubating coverslips with 1X Click-iT OPP Reaction Cocktail that includes 1X Click-iT OPP Reaction Buffer, 1X Copper Protectant, 1X Alexafluor picolyl azide, and 1X Click-iT Reaction Buffer Additive for 30 minutes protected from light. This was removed and Reaction Rinse Buffer was added for 5 minutes. This was removed and coverslips were washed twice with PBS, allowed to dry fully, and mounted with Fluormount-G (Invitrogen, 00-4958-02).

Cells were then imaged on a Nikon Eclipse Ti-E microscope with a Yokogawa, two-camera, CSU0W1 spinning disk system with a Nikon LU-N4 laser launch at room temperature with a 100X oil-immersion objective (1.45 numerical aperture). Images were acquired using Nikon Elements and analyzed using ImageJ as follows. Z-stacks were obtained then combined into sum slices for quantification of maximum intensity projections for viewing. In the summed images, the integrated density and area of individual cells was obtained, as well as the background fluorescence. These were then used to calculate CTCF, which was then normalized to the “Pre” sample for each cell.

### Phalloidin staining and quantification

Procedure adapted from (Craig et al., 2019). Cells were mounted onto poly-lysine coverslips and fixed with fresh 4% paraformaldehyde in 1X HEPES. Coverslips with cells were then permeabilized with acetone and allowed to dry. Cells were rehydrated with PBS, stained with Phalloidin-Atto 488 (Sigma, 49409-10NMOL), and finally washed with PBS and allowed to dry before mounting with Fluormount-G (Craig et al., 2019). Cells were imaged using the Nikon Spinning Disk Confocal discussed above. Z-stacks were obtained in Nikon Elements, and in ImageJ, maximum intensity projections were created for viewing. Publication quality images were acquired using a Zeiss LSM880 with Airyscan with two photomultiplier tubes, a GaAsP detector, and a transmitted light detector. Images were taken at room temperature using a 100x (1.46 numerical aperture) oil-immersion lens. Images were acquired using Zeiss Zen (black edition) and prepared for publication using ImageJ.

### Electron microscopy

Cells (1mL of each strain) were deciliated via pH shock by adding 115μL of 0.5N acetic acid for 45 seconds followed by 120μL of 0.5N KOH to bring cells back to neutral pH. Cells were allowed to regrow cilia for 30 minutes. A pre sample and a 30-minute post-deciliation sample were fixed in an equal volume of 2% glutaraldehyde for 20 minutes at room temperature. Samples were then pelleted using gentle centrifugation for 10 minutes. The supernatant was removed, and cells were resuspended in 1% glutaraldehyde, 20mM sodium cacodylate. Cells were incubated for 1 hour at room temperature and then overnight at 4º Celsius. This protocol was first reported in (Dentler and Adams, 1992). A JEOL JEM-1400 Transmission Electron Microscope equipped with a Lab6 gun was used to acquire images. Images were quantified in ImageJ.

### SAG1-HA Immunofluorescence

Procedure adapted from (Belzile et al., 2013). SAG1-HA cells were grown overnight in M-N media to induce gametes. These cells were then treated with either 10μM LatB for 1 hour or 250μM CK-666 for 2 hours. Following treatment, mating was induced by adding db-cAMP (ChemCruz, Santa Cruz, CA) to a final concentration of 13.5mM and incubating for 30 minutes. Cells were adhered to coverslips and fixed with methanol. Cells were then dried and rehydrated with PBS and incubated with 100% block (5% BSA, 1% fish gelatin) for 30 minutes. The 100% block was replaced with new 100% block containing 10% normal goat serum for another 30-minute incubation. The rabbit anti-HA primary antibody (Cell Signaling, C29F4) was diluted 1:1000 in 20% block in PBS. Coverslips were incubated at 4º Celsius in a humidified chamber overnight. The primary antibody was removed and washed away with 3 10-minute PBS washes. The Alexafluor 488-conjugated goat anti-rabbit secondary (Invitrogen, A-10088) was added and coverslips were incubated at room temperature for 1 hour. This was followed by 3 more 10-minute PBS washes and finally mounting with Fluoromount-G. Cells were imaged using the Nikon Spinning Disk Confocal microscope, lens, and software discussed previously. Z-stacks were obtained, and maximum intensity projections were created for visualization and sum slices were created for quantification using ImageJ.

Images were quantified by using line scans from the apex of the cells to the basal region of the cells farthest away from the apex. Line scans were then normalized, and background subtracted before being combined into single graphs. Using the line scans, the intensity of signal at the basal region of the cells was subtracted from the signal at the apical region. Finally, cells with a difference over 30 were considered to be apically enriched and this was quantified as percentage of cells with apical staining.

### SAG1-HA western blot

Procedure adapted from (Belzile et al., 2013). SAG1-HA cells were grown overnight in M-N media to induce gametes. These cells were then treated with either 10μM LatB for 1 hour or 250μM CK-666 for 2 hours. Following treatment, mating induction was done by adding db-cAMP (ChemCruz, SC-201567B) to a final concentration of 13.5mM and incubating for 10 minutes. Cells were then treated with 0.01% trypsin (Sigma, T8003) for 5 minutes, pelleted (at 500xg for 2 minutes), resuspended in lysis buffer (5% glycerol, 1% NP-40, 1mM DTT, 1X protease inhibitors), and then lysed with bead beating. A western blot was carried out as described above using rabbit anti-HA primary antibody (Cell Signaling, C29F4) diluted to 1:1000 in 1% BSA, 1% milk and HRP-conjugated goat anti-rabbit secondary (Thermo, G-21234) diluted to 1:5000 in 1% milk 1% BSA.

### Chlamydomonas mating

SAG1-HA (mating type plus) and *arpc4* mutants (mating type minus) were incubated in M-N media (minimal media without nitrogen) for 8 hours to induce gamete formation. The two liquid cultures were then mixed and allowed to incubate under white light without agitation overnight. The next day the pellicle was transferred to a 4% TAP plate. This was incubated under white light overnight, then covered in foil and placed in a dark drawer for 5-7 days. After 5-7 days, the zygospores were transferred individually and manually from the 4% TAP plate to a 1.5% TAP plate using a dissecting microscope (Zeiss). This plate was incubated in white light overnight. The following day the zygospores that had split into tetrads were dissected. These were then allowed to grow before being screened via PCR for a colony containing the *arpc4* mutant and SAG1-HA.

### Membrane stain

Cells were treated with either PitStop2 (Sigma, SML1169) or Dynasore hydrate (Sigma, D7693) for 1 hour. Meanwhile, FM 4-64FX membrane stain (Invitrogen, F34653) was diluted to a stock concentration of 200μg/mL. Cells were adhered to poly-lysine coverslips. After a 5-minute incubation, cells were tilted off and 5μg/mL of ice-cold stain in Hank’s Buffered Salt Solution (HBSS) without magnesium or calcium was added for 1 minute. The stain was tilted off and cells were fixed with ice cold 4% paraformaldehyde in HBSS without magnesium or calcium for 15 minutes. Coverslips were then rinsed 3 times for 10 minutes each in ice cold HBSS without magnesium or calcium. Finally, cells were mounted with Fluoromount-G and imaged using the Nikon Spinning Disk Confocal microscope, lens, and software discussed previously. Z-stacks were taken and combined into sum projections using ImageJ. The background corrected total cell fluorescence was then calculated by taking the integrated density and subtracting the sum of the area and the mean background intensity.

### Biotin ciliary isolation

Procedure adapted from (Dentler, 2013). 100mL of cells were grown in TAP for each condition until they reached an OD_730_ of 1.6 or above. Cells were then spun down and resuspended in M1 media and allowed to grow overnight. The next day cells were spun down at 1800rpm for 3 minutes and resuspended in HM Media (10mM HEPES, 5mM MgSO4, pH 7.2). Solid biotin (Thermo, 21335) was added to 20μg/mL for each strain and incubated for 5 minutes with agitation. Cells were diluted with 10 volumes of fresh M1 media before being spun down at 1800rpm for 3 minutes. After all cells were pelleted, they were washed with fresh M1 media three times. A pre sample was set aside (100mL) and the remainder of the cells were resuspended in 4.5 pH M1 media for 45 seconds before being spun down again at 1800rpm for 3 minutes. Cells were then resuspended in pH 7.0 media and allowed to regrow their cilia for 4 hours. A sample was taken pre-biotinylation to use as a control for non-specific streptavidin binding.

Meanwhile, the cilia were isolated from the pre sample. The samples were centrifuged for 3 minutes at 1800rpm. Supernatant was drained and each pellet was resuspended in 2 mL of 10mM HEPES (pH 7.4). This was repeated 2 times. Then each pellet was resuspended in 1 mL of fresh ice-cold 4% HMDS (10mM HEPES pH 7.4, 5mM MgSO4, 1mM DTT, 4% w/v sucrose). Cells were deciliated by incubating with 25mM dibucaine for 2 minutes. Then ice cold HMDS with 0.5mM EGTA was added (1mL per 1.5mL of cells). This was then centrifuged for 3 minutes at 1800rpm. Supernatant was collected for each sample. Then HMDS with 25% sucrose was layered beneath the supernatant (2 mL of 25% HMDS for 1mL of supernatant) to create an interface. This was centrifuged at 4º Celsius for 10 min at 2400rpm with no brake to avoid disrupting interface where cilia should now be located. Cilia were removed, pelleted at 21130xg for 30 minutes, then resuspended in lysis buffer (5% glycerol, 1% NP-40, 1mM DTT, 1X protease inhibitors). This was repeated with the post samples 4 hours following deciliation. Protein gel electrophoresis and blotting was performed as described as above using an HRP-conjugated streptavidin (Thermo, S911).

### Homology modeling and sequence studies

Arp2/3 homology model was created using the Modeller plugin in UCSF Chimera. The template used was 1U2Z (Nolen et al., 2004; Pettersen et al., 2004; Sali and Blundell, 1993). Percent identity and similarity is calculated in relation to the human Arp2/3 complex members using a MUSCLE alignment in Geneious. The homology model was visualized and conservation was mapped on the protein surface using Chimera (Pettersen et al., 2004).

### Statistical analysis

Statistical analyses were done if GraphPad Prism Version 9. Superplots were created using the method in (Lord et al., 2020). For any experiments comparing 2 groups (**Figure 3D, 5C, and 5E**) an unpaired student’s t-test comparing the means of the 3 biological replicates was used to determine P value. For experiments comparing multiple samples (**Figure 1A, 1B, 1C, 1D, 2B, 2C, 3B, and 6E**), an ANOVA was used comparing the means of the 3 biological replicates. This was followed by a multiple comparisons test (Tukey’s). For any percentages shown (**Figure 7D**), Chi-squared analysis was performed. For all experiments **** P<0.0001, *** P<0.001, ** P<0.01, * P<0.1 with p values listed in the figure legends.

## ACKNOWLEDGEMENTS

Our gratitude to William Dentler for providing expertise especially in looking at the electron microscopy images and helpful advice, William Snell for providing the SAG1-HA strain, Masayuki Onishi for the *nap1* strain, Henry Higgs for his feedback on version 1 of the manuscript, Ann Lavanway for assistance with microscopy, and the Avasthi lab for their help throughout the project. We also thank David Sept and Courtney M Schroeder for the help with the original version of this paper and for providing helpful comments.

We thank our funding sources including the Madison and Lila Self Graduate Fellowship at the University of Kansas Medical Center and the MIRA (R35GM128702). Finally, we thank the BioMT core at Dartmouth College (NIH/NIGMS COBRE award P20-GM113132), the Genomics and Molecular Biology Shared Resources Core (NCI Cancer Center Support Grant 5P30CA023108-37), and the KIDDRC NIH U54 HD 090216 at the University of Kansas Medical Center, Kansas City, KS 66160.

The authors have no additional competing financial interests.

## AUTHOR CONTRIBUTIONS

Brae M Bigge: Conceptualization, data curation, formal analysis, investigation, methodology, visualization, writing (original draft), writing (review & editing) Nicholas E Rosenthal: Data curation, formal analysis, writing (review & editing) Prachee Avasthi: Conceptualization, funding acquisition, project administration, resources, supervision

## SUPPLEMENTAL MATERIAL

**Figure S1.**
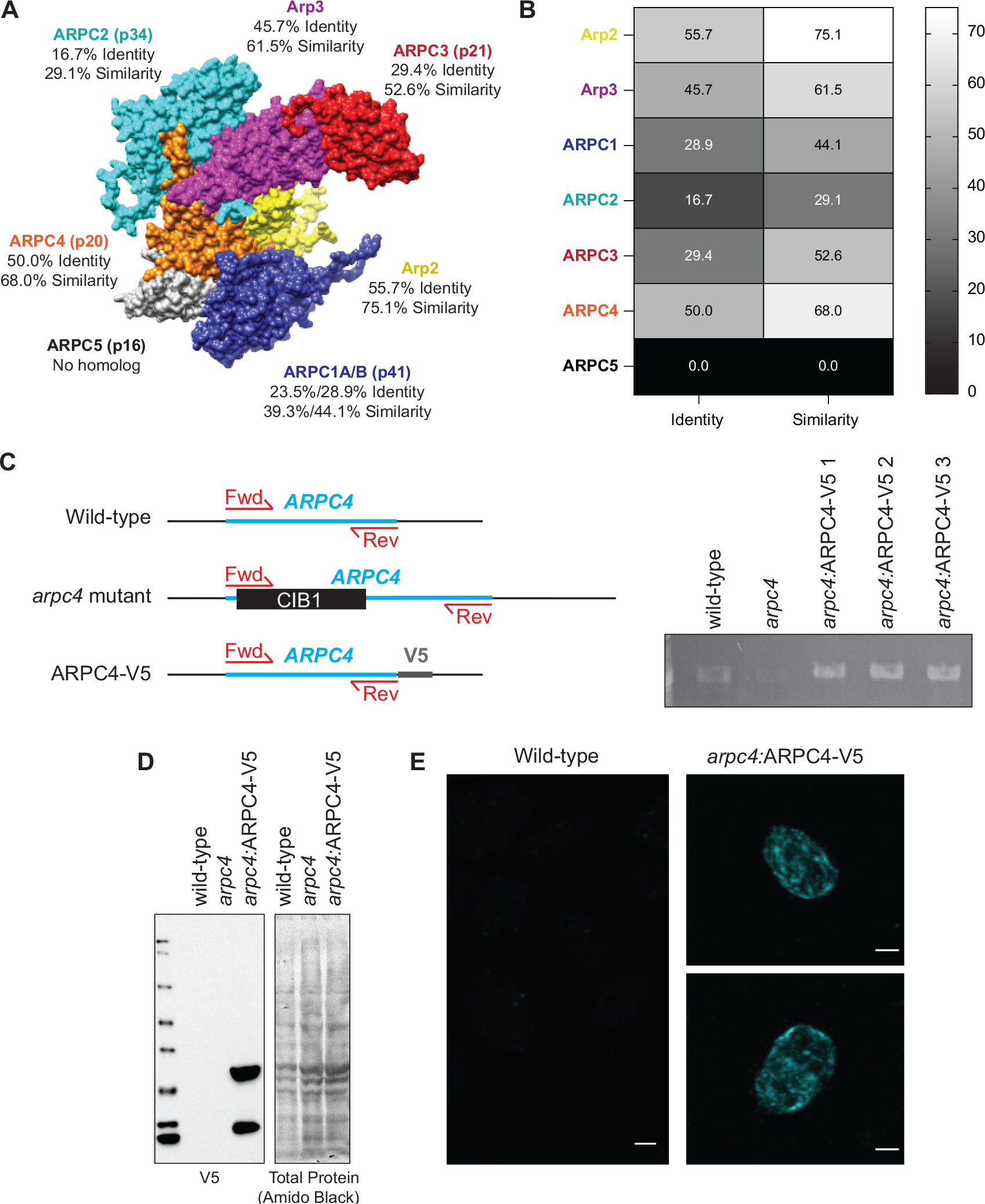
Arp2/3 complex conservation in *Chlamydomonas*. **A)** Homology model of the *Chlamydomonas* Arp2/3 complex based on the bovine Arp2/3 complex (PDB:1K8K). Percent identity and similarity for the protein sequences of the Arp2/3 complex of *Chlamydomonas* compared to the bovine Arp2/3 complex. **B)** Heatmap of sequence identity and similarity of the Arp2/3 complex members of *Chlamydomonas* compared to those of the bovine complex. The ARPC1 isoform used for comparison was ARPC1B as it was more highly conserved to the *Chlamydomonas* ARPC1. Percentages were determined based on a MUSCLE alignment in Geneious. **C)** Diagram of wild-type ARPC4, mutated ARPC4, and ARPC4-V5 with primer position. PCR gel showing presences of the *ARPC4* gene in wild-type and rescue colonies, but not in the *arpc4* mutant. **D)** Western blot using V5 antibody (Thermo) showing protein expression of V5 in rescues containing *ARPC4-V5*. Total protein was probed using amido black. **E)** Immunofluorescence using the V5 antibody (Thermo). Wild-type cells show little to no signal, while cells expressing *ARPC4-V5* on the *arpc4* mutant background (colony 3) do show diffuse signal, suggesting the ARPC4-V5 is present. Scale bar represents 2μm.

**Figure S2.**
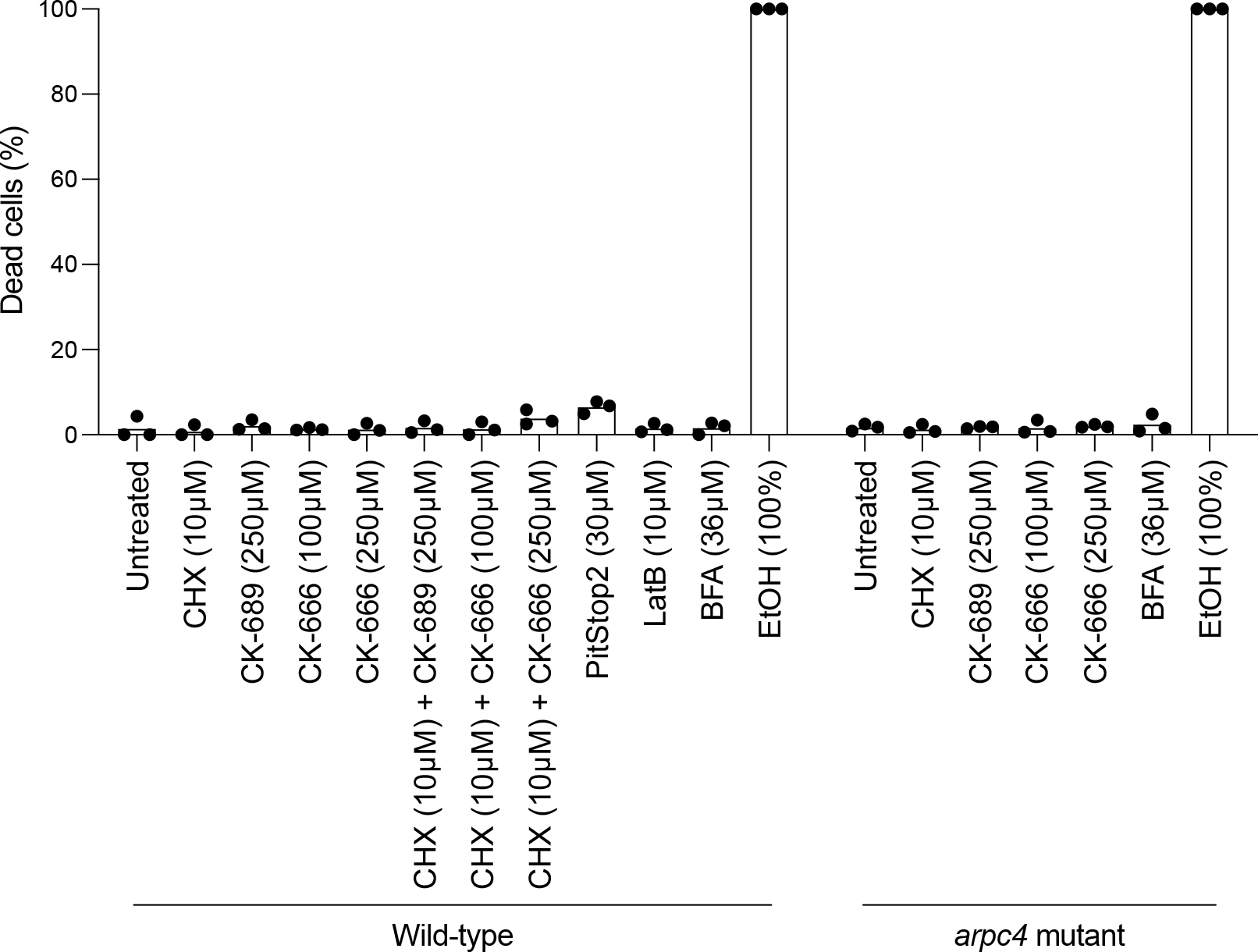
Health of cells treated with chemical inhibitors. For each chemical inhibitor throughout the paper cells were stained with sytox to determine health of the cells. Ethanol (EtOH) is used as a positive control as it kills the cells. Cells were treated with LatB or PitStop2 for 1 hour consistent with what was used in the paper. Cells treated with any concentrations of CK-666, CK-689, or CHX were treated for 2 hours consistent with what was used in the paper and when ciliary growth should be complete. Cells treated with BFA were treated for 3 hours consistent with what was used in the paper. n > 70 cells in 3 separate experiments.

**Figure S3.**
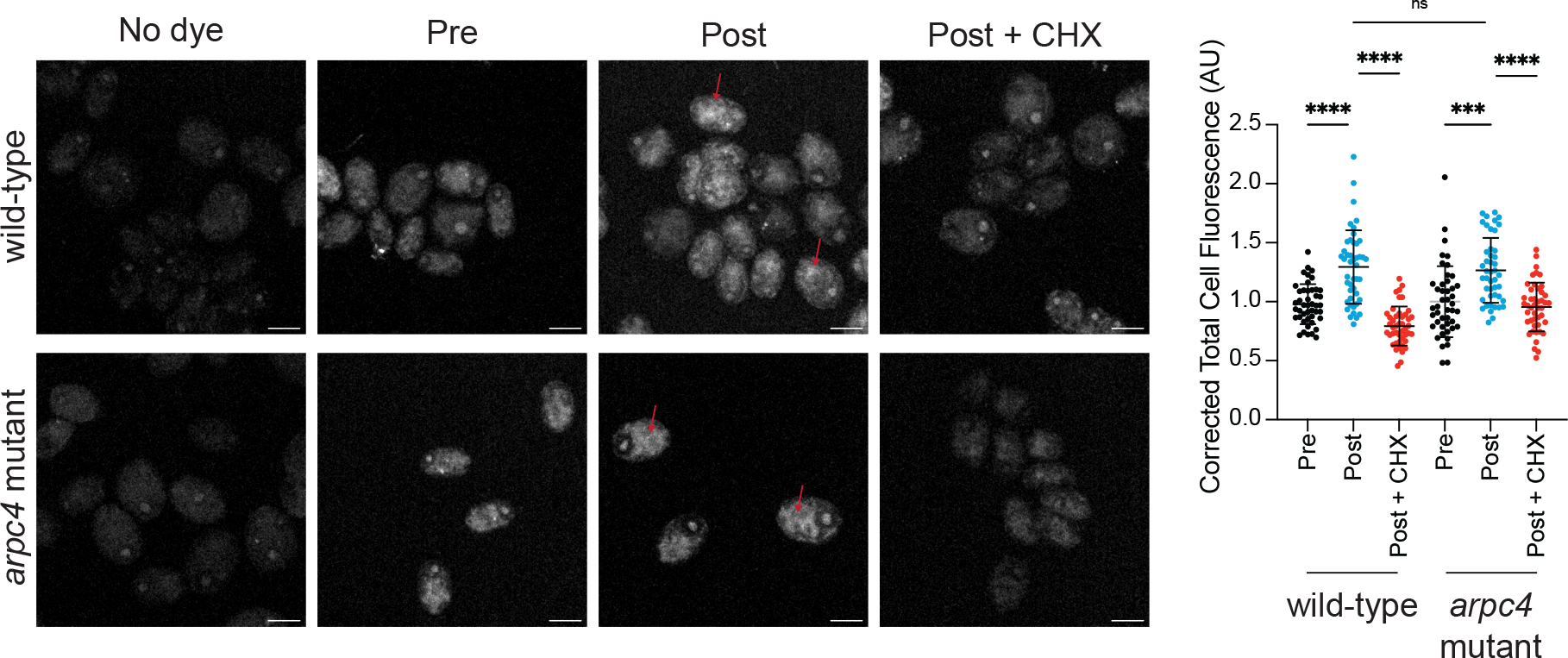
Protein synthesis following deciliation is not defective in *arpc4* mutants. Wild-type and *arpc4* mutant cells were treated with Click-iT OPP either before deciliation, after deciliation and one hour of regrowth, or after deciliation and one hour of regrowth in 10μM CHX which blocks protein translation. Following deciliation there was an increase in fluorescence in cells, particularly around the nucleus (red arrows). The total cell fluorescence was measured and corrected to background then quantified in the graph. n=30 cells per treatment group in 3 separate experiments. **** means P<0.0001. Scale bar represents 5μm.

**Figure S4.**
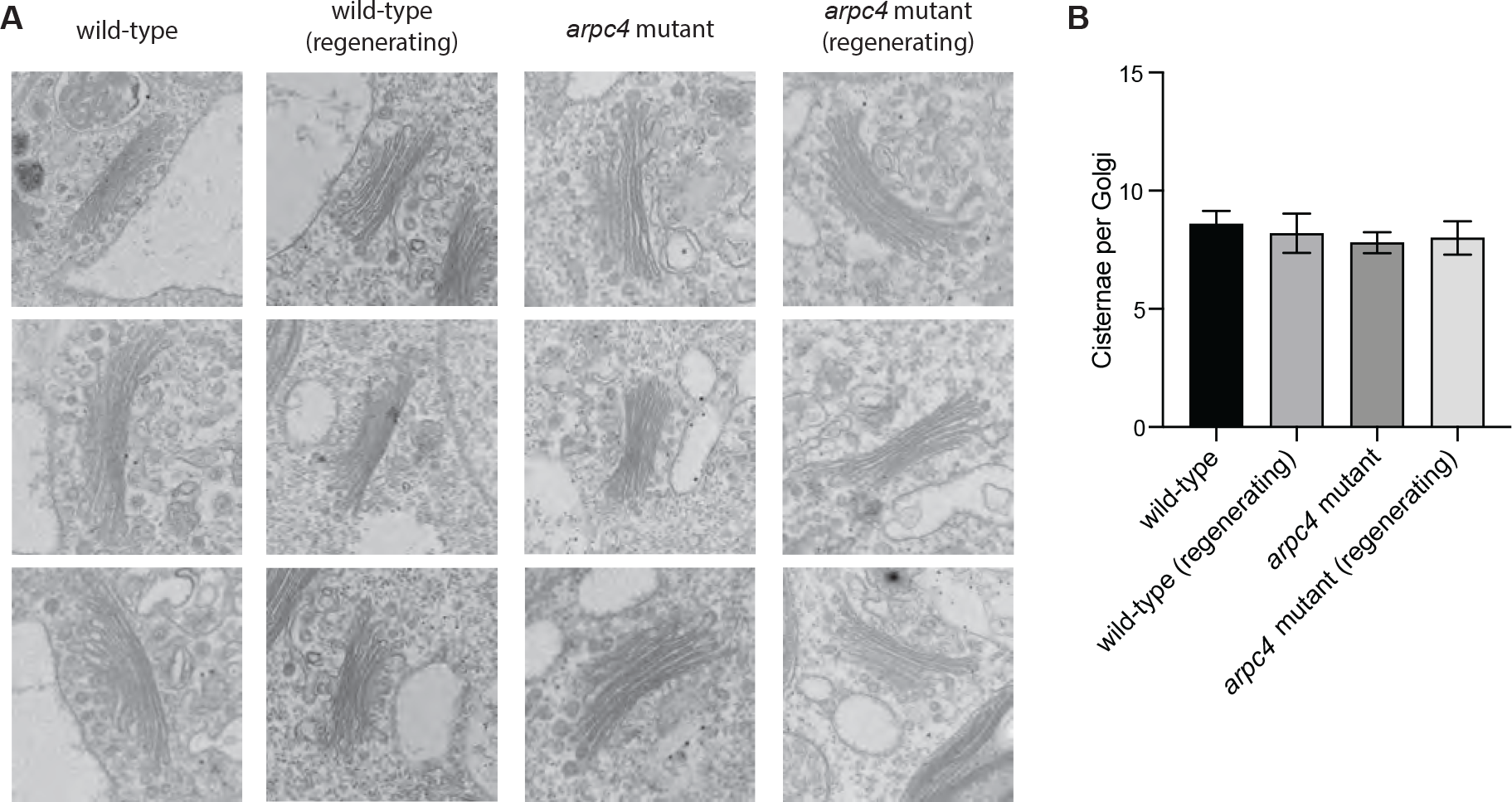
The Arp2/3 complex is not required for Golgi organization. **A)** Transmission electron micrographs of Golgi found in wild-type or *arpc4* cells. **B)** Number of cisternae per Golgi for each condition. n=5. Error bars represent standard deviation.

**Figure S5.**
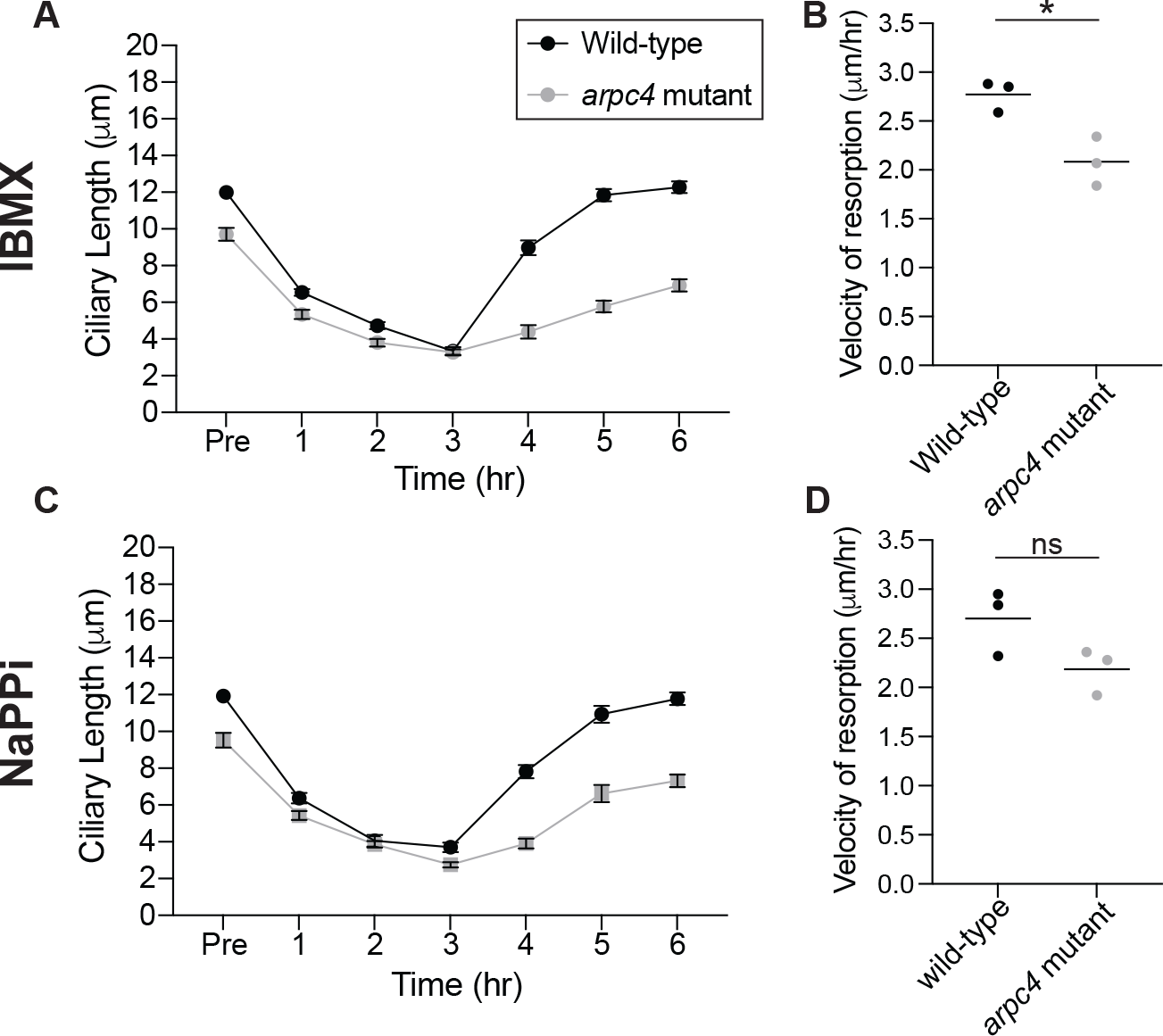
Resorption of cilia with NaPPi and IBMX is not increased in the *arpc4* mutant as it is with BFA. **A)** Cells were treated with 1mM IBMX and allowed to resorb their cilia. After 3 hours, IBMX was washed out and cells were allowed to regrow cilia. n=30 in 3 separate experiments. **B)** The velocity of IBMX resorption was determined by fitting a line to the first 4 points during regeneration and determining the slope in 3 separate experiments. P=0.0158. **C)** Cells were treated with 20mM NaPPi and allowed to resorb their cilia. After 3 hours, NaPPi was washed out and cilia were allowed to regrow. n=30 in 3 separate experiments. **D)** The velocity of NaPPi resorption was determined by fitting a line to the first 4 points of resorption and determining the slope in 3 separate experiments. P=0.0945. The slightly slower velocities of resorption in the *arpc4* mutant may be due to the fact that these cells start with shorter cilia and therefore have less to resorb or it may be due to problems in endocytosis that is thought to be required for resorption of cilia (Saito et al., 2017).

## Notes

### Competing Interest Statement

PA is a paid consultant for Arcadia Science

### Summary of Updates

The paper was reorganized to better highlight the main points of the paper. We also added text throughout to address comments from reviewers and to clarify some of the conclusions. To figure 1, we added arpc4 mutants treated with CK-666 to demonstrate the specificity of CK-666 in this organism. This was previously in the supplement, but we felt that it should be moved to the main figure. The previous figure 5 (now figure 7) was updated. The clathrin light chain staining was removed as we felt it was likely that is antibody had non-specific targets and therefore may have been confounding the data. To this figure we also included another version of the membrane dye experiment this time treating cells with CK-666 to show that the phenotype was conserved whether we blocked the Arp2/3 complex with a genetic mutation or a chemical inhibitor. To previous figure 8 (now figure 4) we added no biotin controls. We also added a diagram to supplemental figure 2 showing the sequence of the ARPC4 gene and how it was affected by the CIB cassette.

## REFERENCES

Adams, A.E., Pringle, J.R., 1984. Relationship of actin and tubulin distribution to bud growth in wild-type and morphogenetic-mutant Saccharomyces cerevisiae. J. Cell Biol. 98, 934– 945. https://doi.org/10.1083/jcb.98.3.934

Aghamohammadzadeh, S., Ayscough, K.R., 2009. Differential requirements for actin during yeast and mammalian endocytosis. Nat. Cell Biol. 11, 1039–1042. https://doi.org/10.1038/ncb1918

Avasthi, P., Onishi, M., Karpiak, J., Yamamoto, R., Mackinder, L., Jonikas, M.C., Sale, W.S., Shoichet, B., Pringle, J.R., Marshall, W.F., 2014. Actin Is Required for IFT Regulation in Chlamydomonas reinhardtii. Curr. Biol. 24, 2025–2032. https://doi.org/10.1016/j.cub.2014.07.038

Ayscough, K.R., Stryker, J., Pokala, N., Sanders, M., Crews, P., Drubin, D.G., 1997. High rates of actin filament turnover in budding yeast and roles for actin in establishment and maintenance of cell polarity revealed using the actin inhibitor latrunculin-A. J. Cell Biol. 137, 399–416. https://doi.org/10.1083/jcb.137.2.399

Basu, R., Munteanu, E.L., Chang, F., 2014. Role of turgor pressure in endocytosis in fission yeast. Mol. Biol. Cell 25, 679–687. https://doi.org/10.1091/mbc.E13-10-0618

Belzile, O., Hernandez-Lara, C.I., Wang, Q., Snell, W.J., 2013. Regulated membrane protein entry into flagella is facilitated by cytoplasmic microtubules and does not require IFT. Curr. Biol. CB 23, 1460–1465. https://doi.org/10.1016/j.cub.2013.06.025

Bouck, G.B., 1971. THE STRUCTURE, ORIGIN, ISOLATION, AND COMPOSITION OF THE TUBULAR MASTIGONEMES OF THE OCHROMONAS FLAGELLUM. J. Cell Biol. 50, 362–384. https://doi.org/10.1083/jcb.50.2.362

Campellone, K., Welch, M., 2010. A nucleator arms race: cellular control of actin assembly. Nat. Rev. Mol. Cell Biol. 11, 237–251.

Carlsson, A.E., Bayly, P.V., 2014. Force generation by endocytic actin patches in budding yeast. Biophys. J. 106, 1596–1606. https://doi.org/10.1016/j.bpj.2014.02.035

Cheng, X., Liu, G., Ke, W., Zhao, L., Lv, B., Ma, X., Xu, N., Xia, X., Deng, X., Zheng, C., Huang, K., 2017. Building a multipurpose insertional mutant library for forward and reverse genetics in Chlamydomonas. Plant Methods 13, 36. https://doi.org/10.1186/s13007-017-0183-5

Cochilla, A.J., Angleson, J.K., Betz, W.J., 1999. MONITORING SECRETORY MEMBRANE WITH FM1-43 FLUORESCENCE. Annu. Rev. Neurosci. 22, 1–10. https://doi.org/10.1146/annurev.neuro.22.1.1

Craig, E.W., Mueller, D.M., Bigge, B.M., Schaffer, M., Engel, B.D., Avasthi, P., 2019. The elusive actin cytoskeleton of a green alga expressing both conventional and divergent actins. Mol. Biol. Cell mbc.E19-03-0141. https://doi.org/10.1091/mbc.E19-03-0141

Dentler, W., 2013. A Role for the Membrane in Regulating Chlamydomonas Flagellar Length. PLOS ONE 8, e53366. https://doi.org/10.1371/journal.pone.0053366

Dentler, W.L., Adams, C., 1992. Flagellar microtubule dynamics in Chlamydomonas: cytochalasin D induces periods of microtubule shortening and elongation; and colchicine induces disassembly of the distal, but not proximal, half of the flagellum. J. Cell Biol. 117, 1289–1298. https://doi.org/10.1083/jcb.117.6.1289

Diener, D.R., Lupetti, P., Rosenbaum, J.L., 2015. Proteomic analysis of isolated ciliary transition zones reveals the presence of ESCRT proteins. Curr. Biol. CB 25, 379–384. https://doi.org/10.1016/j.cub.2014.11.066

Farina, F., Gaillard, J., Guérin, C., Couté, Y., Sillibourne, J., Blanchoin, L., Théry, M., 2016. The centrosome is an actin-organizing centre. Nat. Cell Biol. 18, 65–75. https://doi.org/10.1038/ncb3285

Gachet, Y., Hyams, J.S., 2005. Endocytosis in fission yeast is spatially associated with the actin cytoskeleton during polarised cell growth and cytokinesis. J. Cell Sci. 118, 4231–4242. https://doi.org/10.1242/jcs.02530

Goode, B.L., Eskin, J.A., Wendland, B., 2015. Actin and endocytosis in budding yeast. Genetics 199, 315–358. https://doi.org/10.1534/genetics.112.145540

Gournier, H., Goley, E.D., Niederstrasser, H., Trinh, T., Welch, M.D., 2001. Reconstitution of Human Arp2/3 Complex Reveals Critical Roles of Individual Subunits in Complex Structure and Activity. Mol. Cell 8, 1041–1052. https://doi.org/10.1016/S1097-2765(01)00393-8

Hetrick, B., Han, M.S., Helgeson, L.A., Nolen, B.J., 2013. Small Molecules CK-666 and CK-869 Inhibit Actin-Related Protein 2/3 Complex by Blocking an Activating Conformational Change. Chem. Biol. 20, 701–712. https://doi.org/10.1016/j.chembiol.2013.03.019

Hirono, M., Uryu, S., Ohara, A., Kato-Minoura, T., Kamiya, R., 2003. Expression of conventional and unconventional actins in Chlamydomonas reinhardtii upon deflagellation and sexual adhesion. Eukaryot. Cell 2, 486–493. https://doi.org/10.1128/ec.2.3.486-493.2003

Hu Qicong, Milenkovic Ljiljana, Jin Hua, Scott Matthew P., Nachury Maxence V., Spiliotis Elias T., Nelson W. James, 2010. A Septin Diffusion Barrier at the Base of the Primary Cilium Maintains Ciliary Membrane Protein Distribution. Science 329, 436–439. https://doi.org/10.1126/science.1191054

Inoue, D., Obino, D., Pineau, J., Farina, F., Gaillard, J., Guerin, C., Blanchoin, L., Lennon-Duménil, A.-M., Théry, M., 2019. Actin filaments regulate microtubule growth at the centrosome. EMBO J. 38. https://doi.org/10.15252/embj.201899630

Jack, B., Avasthi, P., 2018. Erratum to: Chemical Screening for Flagella-Associated Phenotypes in Chlamydomonas reinhardtii. Methods Mol. Biol. Clifton NJ 1795, E1. https://doi.org/10.1007/978-1-4939-7874-8_19

Jack, B., Mueller, D.M., Fee, A.C., Tetlow, A.L., Avasthi, P., 2019. Partially Redundant Actin Genes in Chlamydomonas Control Transition Zone Organization and Flagellum-Directed Traffic. Cell Rep. 27, 2459-2467.e3. https://doi.org/10.1016/j.celrep.2019.04.087

Jin, H., White, S.R., Shida, T., Schulz, S., Aguiar, M., Gygi, S.P., Bazan, J.F., Nachury, M.V., 2010. The conserved Bardet-Biedl syndrome proteins assemble a coat that traffics membrane proteins to cilia. Cell 141, 1208–1219. https://doi.org/10.1016/j.cell.2010.05.015

Kaplan, O.I., Doroquez, D.B., Cevik, S., Bowie, R.V., Clarke, L., Sanders, A.A.W.M., Kida, K., Rappoport, J.Z., Sengupta, P., Blacque, O.E., 2012. Endocytosis Genes Facilitate Protein and Membrane Transport in C. elegans Sensory Cilia. Curr. Biol. 22, 451–460. https://doi.org/10.1016/j.cub.2012.01.060

Kato-Minoura, T., Uryu, S., Hirono, M., Kamiya, R., 1998. Highly divergent actin expressed in a Chlamydomonas mutant lacking the conventional actin gene. Biochem. Biophys. Res. Commun. 251, 71–76. https://doi.org/10.1006/bbrc.1998.9373

Kiesel, P., Alvarez Viar, G., Tsoy, N., Maraspini, R., Gorilak, P., Varga, V., Honigmann, A., Pigino, G., 2020. The molecular structure of mammalian primary cilia revealed by cryo-electron tomography. Nat. Struct. Mol. Biol. https://doi.org/10.1038/s41594-020-0507-4

Kim, J., Lee, J.E., Heynen-Genel, S., Suyama, E., Ono, K., Lee, K., Ideker, T., Aza-Blanc, P., Gleeson, J.G., 2010. Functional genomic screen for modulators of ciliogenesis and cilium length. Nature 464, 1048–1051. https://doi.org/10.1038/nature08895

Langousis, G., Shimogawa, M.M., Saada, E.A., Vashisht, A.A., Spreafico, R., Nager, A.R., Barshop, W.D., Nachury, M.V., Wohlschlegel, J.A., Hill, K.L., 2016. Loss of the BBSome perturbs endocytic trafficking and disrupts virulence of Trypanosoma brucei. Proc. Natl. Acad. Sci. U. S. A. 113, 632–637. https://doi.org/10.1073/pnas.1518079113

Lefebvre, P.A., 1995. Flagellar amputation and regeneration in Chlamydomonas, in: Methods in Cell Biology. Elsevier, pp. 3–7.

Lefebvre, P.A., Nordstrom, S.A., Moulder, J.E., Rosenbaum, J.L., 1978. Flagellar elongation and shortening in Chlamydomonas. IV. Effects of flagellar detachment, regeneration, and resorption on the induction of flagellar protein synthesis. J. Cell Biol. 78, 8–27. https://doi.org/10.1083/jcb.78.1.8

Li, X., Patena, W., Fauser, F., Jinkerson, R.E., Saroussi, S., Meyer, M.T., Ivanova, N., Robertson, J.M., Yue, R., Zhang, R., Vilarrasa-Blasi, J., Wittkopp, T.M., Ramundo, S., Blum, S.R., Goh, A., Laudon, M., Srikumar, T., Lefebvre, P.A., Grossman, A.R., Jonikas, M.C., 2019. A genome-wide algal mutant library and functional screen identifies genes required for eukaryotic photosynthesis. Nat. Genet. 51, 627–635. https://doi.org/10.1038/s41588-019-0370-6

Lord, S.J., Velle, K.B., Mullins, R.D., Fritz-Laylin, L.K., 2020. SuperPlots: Communicating reproducibility and variability in cell biology. J. Cell Biol. 219. https://doi.org/10.1083/jcb.202001064

Macia, E., Ehrlich, M., Massol, R., Boucrot, E., Brunner, C., Kirchhausen, T., 2006. Dynasore, a cell-permeable inhibitor of dynamin. Dev. Cell 10, 839–850. https://doi.org/10.1016/j.devcel.2006.04.002

Milenkovic, L., Scott, M.P., Rohatgi, R., 2009. Lateral transport of Smoothened from the plasma membrane to the membrane of the cilium. J. Cell Biol. 187, 365–374. https://doi.org/10.1083/jcb.200907126

Molla-Herman, A., Ghossoub, R., Blisnick, T., Meunier, A., Serres, C., Silbermann, F., Emmerson, C., Romeo, K., Bourdoncle, P., Schmitt, A., Saunier, S., Spassky, N., Bastin, P., Benmerah, A., 2010. The ciliary pocket: an endocytic membrane domain at the base of primary and motile cilia. J. Cell Sci. 123, 1785–1795. https://doi.org/10.1242/jcs.059519

Monis, W.J., Faundez, V., Pazour, G.J., 2017. BLOC-1 is required for selective membrane protein trafficking from endosomes to primary cilia. J. Cell Biol. 216, 2131–2150. https://doi.org/10.1083/jcb.201611138

Mooren Olivia L., Schafer Dorothy A., 2009. Constricting membranes at the nano and micro scale. Proc. Natl. Acad. Sci. 106, 20559–20560. https://doi.org/10.1073/pnas.0911630106

Nachury, M.V., Loktev, A.V., Zhang, Q., Westlake, C.J., Peränen, J., Merdes, A., Slusarski, D.C., Scheller, R.H., Bazan, J.F., Sheffield, V.C., Jackson, P.K., 2007. A Core Complex of BBS Proteins Cooperates with the GTPase Rab8 to Promote Ciliary Membrane Biogenesis. Cell 129, 1201–1213. https://doi.org/10.1016/j.cell.2007.03.053

Nachury, M.V., Seeley, E.S., Jin, H., 2010. Trafficking to the ciliary membrane: how to get across the periciliary diffusion barrier? Annu. Rev. Cell Dev. Biol. 26, 59–87. https://doi.org/10.1146/annurev.cellbio.042308.113337

Nolen, B.J., Littlefield, R.S., Pollard, T.D., 2004. Crystal structures of actin-related protein 2/3 complex with bound ATP or ADP. Proc. Natl. Acad. Sci. U. S. A. 101, 15627. https://doi.org/10.1073/pnas.0407149101

Onishi, M., Pecani, K., Jones, T. th, Pringle, J.R., Cross, F.R., 2018. F-actin homeostasis through transcriptional regulation and proteasome-mediated proteolysis. Proc Natl Acad Sci U A 115, E6487–e6496. https://doi.org/10.1073/pnas.1721935115

Onishi, M., Pringle, J.R., Cross, F.R., 2016. Evidence That an Unconventional Actin Can Provide Essential F-Actin Function and That a Surveillance System Monitors F-Actin Integrity in Chlamydomonas. Genetics 202, 977–96. https://doi.org/10.1534/genetics.115.184663

Onishi, M., Umen, J.G., Cross, F.R., Pringle, J.R., 2019. Cleavage-furrow formation without F-actin in <em>Chlamydomonas</em>. bioRxiv 789016. https://doi.org/10.1101/789016

Papermaster, D.S., Schneider, B.G., Besharse, J.C., 1985. Vesicular transport of newly synthesized opsin from the Golgi apparatus toward the rod outer segment. Ultrastructural immunocytochemical and autoradiographic evidence in Xenopus retinas. Invest. Ophthalmol. Vis. Sci. 26, 1386–1404.

Park, R.J., Shen, H., Liu, L., Liu, X., Ferguson, S.M., De Camilli, P., 2013. Dynamin triple knockout cells reveal off target effects of commonly used dynamin inhibitors. J. Cell Sci. 126, 5305–5312. https://doi.org/10.1242/jcs.138578

Park, T.J., Mitchell, B.J., Abitua, P.B., Kintner, C., Wallingford, J.B., 2008. Dishevelled controls apical docking and planar polarization of basal bodies in ciliated epithelial cells. Nat. Genet. 40, 871–879. https://doi.org/10.1038/ng.104

Pasquale, S.M., Goodenough, U.W., 1987. Cyclic AMP functions as a primary sexual signal in gametes of Chlamydomonas reinhardtii. J. Cell Biol. 105, 2279–2292. https://doi.org/10.1083/jcb.105.5.2279

Pedersen, L.B., Rosenbaum, J.L., 2008. Intraflagellar transport (IFT) role in ciliary assembly, resorption and signalling. Curr. Top. Dev. Biol. 85, 23–61. https://doi.org/10.1016/S0070-2153(08)00802-8

Pettersen, E.F., Goddard, T.D., Huang, C.C., Couch, G.S., Greenblatt, D.M., Meng, E.C., Ferrin, T.E., 2004. UCSF Chimera--a visualization system for exploratory research and analysis. J. Comput. Chem. 25, 1605–1612. https://doi.org/10.1002/jcc.20084

Ranjan, P., Awasthi, M., Snell, W.J., 2019. Transient Internalization and Microtubule-Dependent Trafficking of a Ciliary Signaling Receptor from the Plasma Membrane to the Cilium. Curr. Biol. 29, 2942-2947.e2. https://doi.org/10.1016/j.cub.2019.07.022

Robinson, R.C., Turbedsky, K., Kaiser, D.A., Marchand, J.-B., Higgs, H.N., Choe, S., Pollard, T.D., 2001. Crystal Structure of Arp2/3 Complex. Science 294, 1679. https://doi.org/10.1126/science.1066333

Rohatgi, R., Snell, W.J., 2010. The ciliary membrane. Curr. Opin. Cell Biol. 22, 541–546. https://doi.org/10.1016/j.ceb.2010.03.010

Rosenbaum, J.L., Moulder, J.E., Ringo, D.L., 1969. Flagellar elongation and shortening in Chlamydomonas. The use of cycloheximide and colchicine to study the synthesis and assembly of flagellar proteins. J. Cell Biol. 41, 600–619. https://doi.org/10.1083/jcb.41.2.600

Saito, M., Otsu, W., Hsu, K.-S., Chuang, J.-Z., Yanagisawa, T., Shieh, V., Kaitsuka, T., Wei, F.-Y., Tomizawa, K., Sung, C.-H., 2017. Tctex-1 controls ciliary resorption by regulating branched actin polymerization and endocytosis. EMBO Rep. 18, 1460–1472. https://doi.org/10.15252/embr.201744204

Sali, A., Blundell, T.L., 1993. Comparative protein modelling by satisfaction of spatial restraints. J. Mol. Biol. 234, 779–815. https://doi.org/10.1006/jmbi.1993.1626

Spector, I., Shochet, N.R., Blasberger, D., Kashman, Y., 1989. Latrunculins—novel marine macrolides that disrupt microfilament organization and affect cell growth: I. Comparison with cytochalasin D. Cell Motil. 13, 127–144. https://doi.org/10.1002/cm.970130302

von Loeffelholz, O., Purkiss, A., Cao, L., Kjaer, S., Kogata, N., Romet-Lemonne, G., Way, M., Moores, C.A., 2020. Cryo-EM of human Arp2/3 complexes provides structural insights into actin nucleation modulation by ARPC5 isoforms. bioRxiv 2020.05.01.071704. https://doi.org/10.1101/2020.05.01.071704

Willox, A.K., Sahraoui, Y.M.E., Royle, S.J., 2014. Non-specificity of Pitstop 2 in clathrin-mediated endocytosis. Biol. Open 3, 326–331. https://doi.org/10.1242/bio.20147955

Wingfield, J.L., Mengoni, I., Bomberger, H., Jiang, Y.-Y., Walsh, J.D., Brown, J.M., Picariello, T., Cochran, D.A., Zhu, B., Pan, J., Eggenschwiler, J., Gaertig, J., Witman, G.B., Kner, P., Lechtreck, K., 2017. IFT trains in different stages of assembly queue at the ciliary base for consecutive release into the cilium. eLife 6, e26609. https://doi.org/10.7554/eLife.26609

Winter, D.C., Choe, E.Y., Li, R., 1999. Genetic dissection of the budding yeast Arp2/3 complex: a comparison of the in vivo and structural roles of individual subunits. Proc. Natl. Acad. Sci. U. S. A. 96, 7288–7293. https://doi.org/10.1073/pnas.96.13.7288

Wood, C.R., Rosenbaum, J.L., 2014. Proteins of the Ciliary Axoneme Are Found on Cytoplasmic Membrane Vesicles during Growth of Cilia. Curr. Biol. 24, 1114–1120. https://doi.org/10.1016/j.cub.2014.03.047

Wu, C.-T., Chen, H.-Y., Tang, T.K., 2018. Myosin-Va is required for preciliary vesicle transportation to the mother centriole during ciliogenesis. Nat. Cell Biol. 20, 175–185. https://doi.org/10.1038/s41556-017-0018-7

Yamada, H., Abe, T., Li, S.-A., Masuoka, Y., Isoda, M., Watanabe, M., Nasu, Y., Kumon, H., Asai, A., Takei, K., 2009. Dynasore, a dynamin inhibitor, suppresses lamellipodia formation and cancer cell invasion by destabilizing actin filaments. Biochem. Biophys. Res. Commun. 390, 1142–1148. https://doi.org/10.1016/j.bbrc.2009.10.105

Zuo, X., Guo, W., Lipschutz, J.H., 2009. The Exocyst Protein Sec10 Is Necessary for Primary Ciliogenesis and Cystogenesis In Vitro. Mol. Biol. Cell 20, 2522–2529. https://doi.org/10.1091/mbc.e08-07-0772

