## SUPPLEMENTAL MATERIAL for "Initial ciliary assembly in *Chlamydomonas* requires Arp2/3 complex-dependent endocytosis"

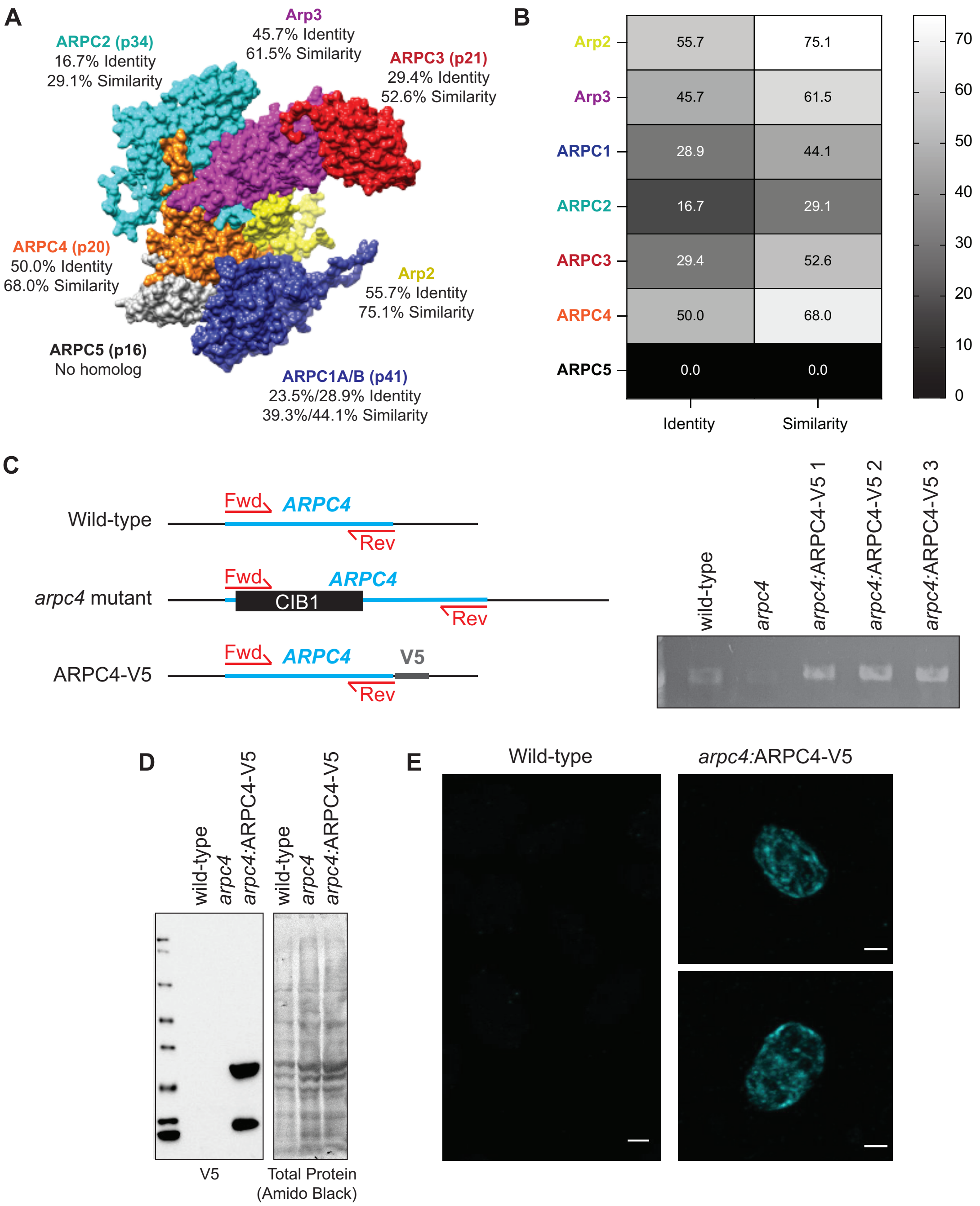


**Figure S1. Arp2/3 complex conservation in *Chlamydomonas*. A)** Homology model of the *Chlamydomonas* Arp2/3 complex based on the bovine Arp2/3 complex (PDB:1K8K). Percent identity and similarity for the protein sequences of the Arp2/3 complex of *Chlamydomonas* compared to the bovine Arp2/3 complex. **B)** Heatmap of sequence identity and similarity of the Arp2/3 complex members of *Chlamydomonas* compared to those of the bovine complex. The ARPC1 isoform used for comparison was ARPC1B as it was more highly conserved to the *Chlamydomonas* ARPC1. Percentages were determined based on a MUSCLE alignment in Geneious. **C)** Diagram of wild-type ARPC4, mutated ARPC4, and ARPC4-V5 with primer position. PCR gel showing presences of the *ARPC4* gene in wild-type and rescue colonies, but not in the *arpc4* mutant. **D)** Western blot using V5 antibody (Thermo) showing protein expression of V5 in rescues containing *ARPC4-V5*. Total protein was probed using amido black. **E)** Immunofluorescence using the V5 antibody (Thermo). Wild-type cells show little to no signal, while cells expressing *ARPC4-V5* on the *arpc4* mutant background (colony 3) do show diffuse signal, suggesting the ARPC4-V5 is present. Scale bar represents 2μm.

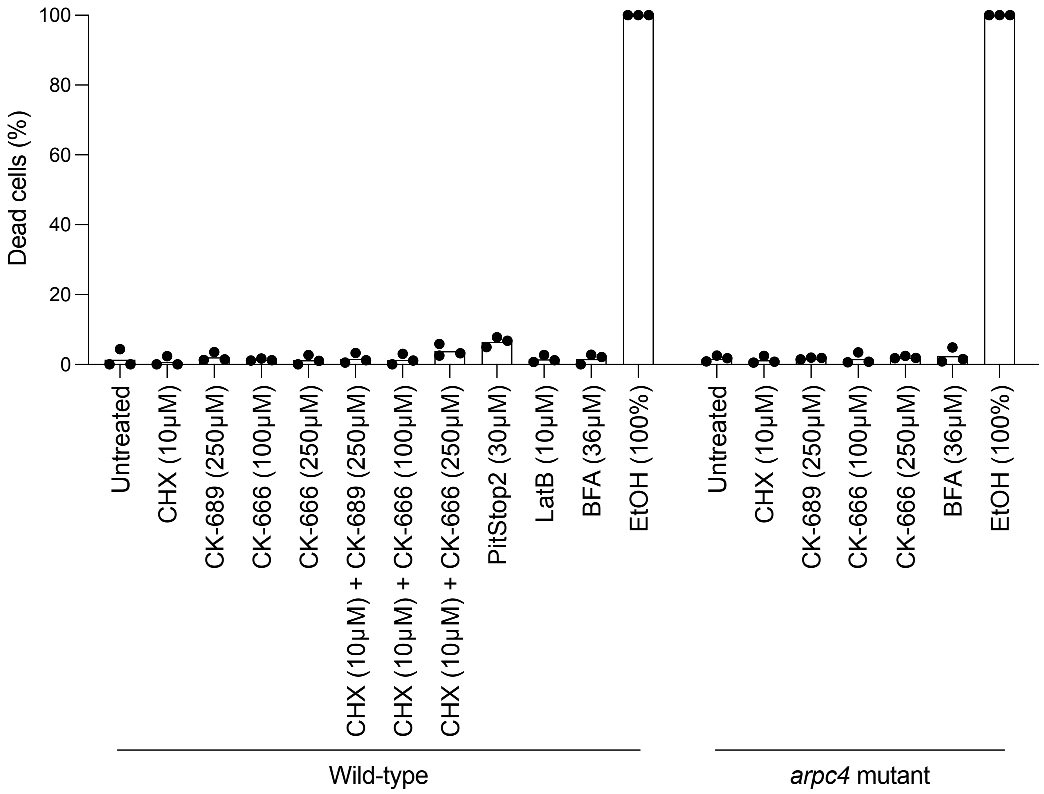


**Figure S2. Health of cells treated with chemical inhibitors.** For each chemical inhibitor throughout the paper cells were stained with sytox to determine health of the cells. Ethanol (EtOH) is used as a positive control as it kills the cells. Cells were treated with LatB or PitStop2 for 1 hour consistent with what was used in the paper. Cells treated with any concentrations of CK-666, CK-689, or CHX were treated for 2 hours consistent with what was used in the paper and when ciliary growth should be complete. Cells treated with BFA were treated for 3 hours consistent with what was used in the paper. n > 70 cells in 3 separate experiments.

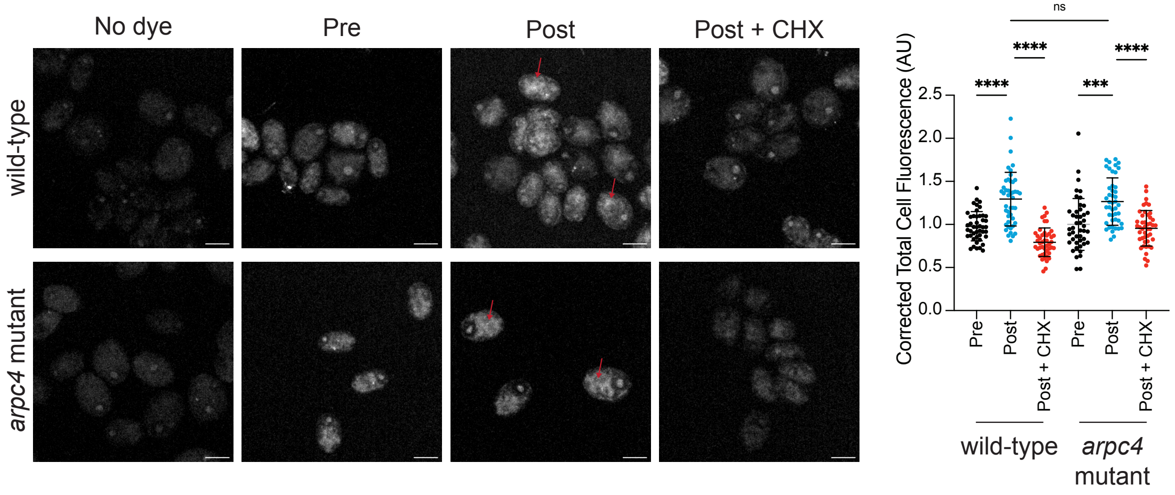


**Figure S3. Protein synthesis following deciliation is not defective in *arpc4* mutants.** Wild-type and *arpc4* mutant cells were treated with Click-iT OPP either before deciliation, after deciliation and one hour of regrowth, or after deciliation and one hour of regrowth in 10µM CHX which blocks protein translation. Following deciliation there was an increase in fluorescence in cells, particularly around the nucleus (red arrows). The total cell fluorescence was measured and corrected to background then quantified in the graph. n=30 cells per treatment group in 3 separate experiments. **** means P<0.0001. Scale bar represents 5µm.

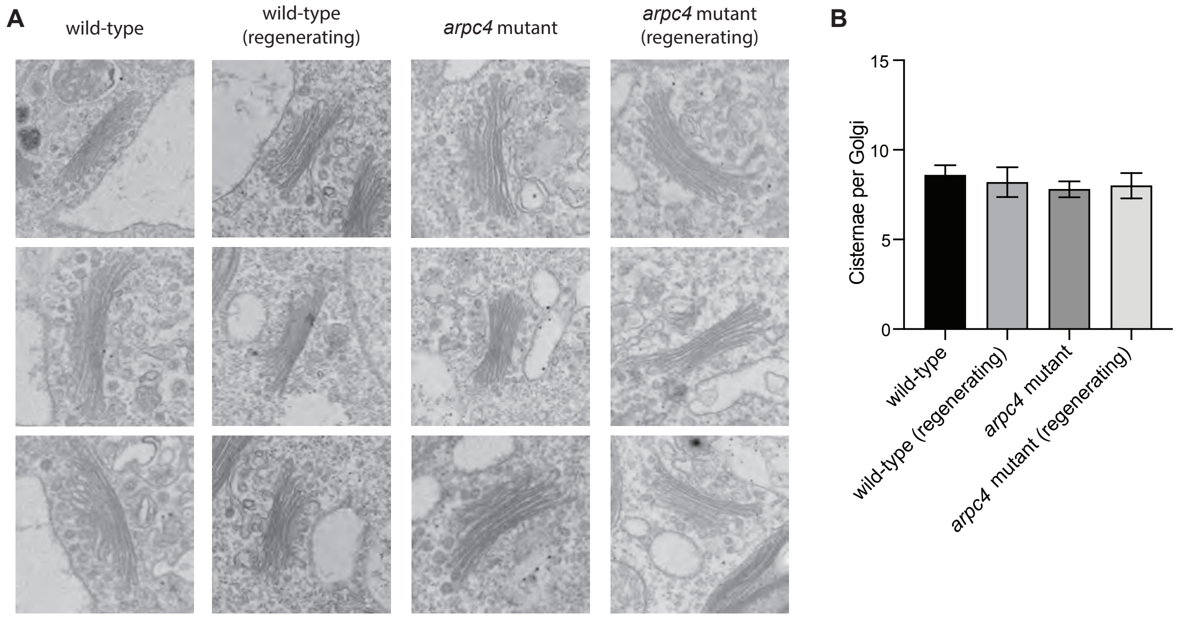


**Figure S4. The Arp2/3 complex is not required for Golgi organization. A)** Transmission electron micrographs of Golgi found in wild-type or *arpc4* cells. **B)** Number of cisternae per Golgi for each condition. n=5. Error bars represent standard deviation.

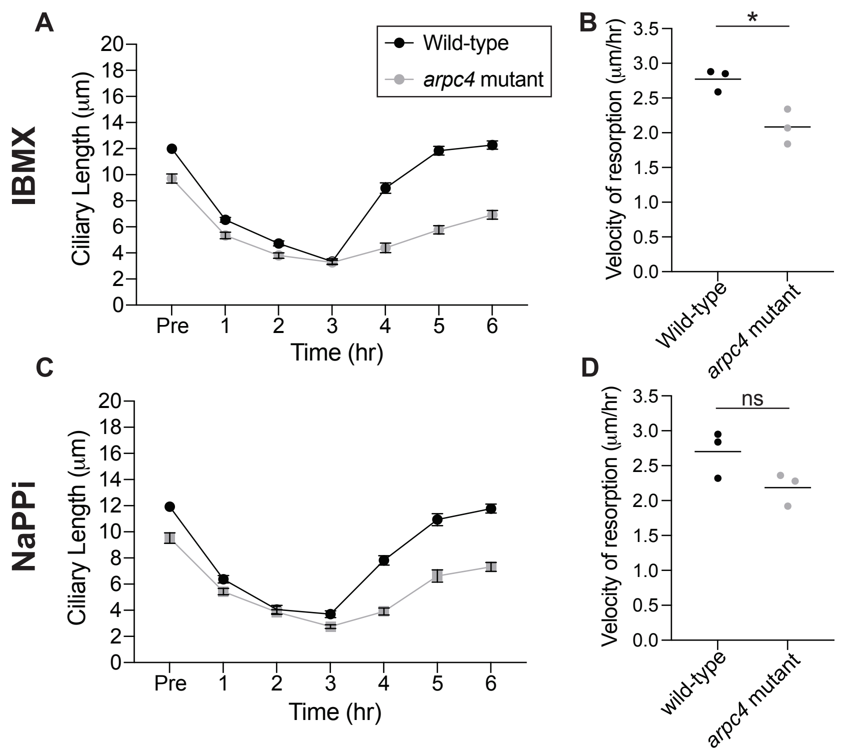


**Figure S5. Resorption of cilia with NaPPi and IBMX is not increased in the *arpc4* mutant as it is with BFA. A)** Cells were treated with 1mM IBMX and allowed to resorb their cilia. After 3 hours, IBMX was washed out and cells were allowed to regrow cilia. n=30 in 3 separate experiments**. B)** The velocity of IBMX resorption was determined by fitting a line to the first 4 points during regeneration and determining the slope in 3 separate experiments. P=0.0158. **C)** Cells were treated with 20mM NaPPi and allowed to resorb their cilia. After 3 hours, NaPPi was washed out and cilia were allowed to regrow. n=30 in 3 separate experiments. **D)** The velocity of NaPPi resorption was determined by fitting a line to the first 4 points of resorption and determining the slope in 3 separate experiments. P=0.0945. The slightly slower velocities of resorption in the *arpc4* mutant may be due to the fact that these cells start with shorter cilia and therefore have less to resorb or it may be due to problems in endocytosis that is thought to be required for resorption of cilia (Saito et al., 2017).
